# A recent shift in centromere size and DNA content in *Podospora pseudocomata* co-occurs with the loss of a fungal genome defense system

**DOI:** 10.64898/2025.12.02.690432

**Authors:** Ivar Westerberg, Mengyuan Li, Eve Mercier, S. Lorena Ament-Velásquez, Linnea Sandell, Philippe Silar, Aaron Vogan, Pierre Grognet, Fabienne Malagnac, Hanna Johannesson

## Abstract

The centromere of the eukaryotic chromosome is necessary for the accurate segregation during cell division. Yet, centromeric DNA is highly variable and rapidly evolving. In fungi, centromeres range from point- to regional centromeres, some of which are hundreds of thousands of base pairs long and filled with transposable elements. As fungi have evolved several specialized defense mechanisms against transposable elements, these regional centromeres are intriguing sites for investigating the connection between genome defense and centromere evolution. Here, we investigated the structure of the centromeres of seven species of the *Podospora anserina* species complex, which is made up of closely related filamentous ascomycetes that diverged less than 1 MYA. We discovered that one species in the complex, *P. pseudocomata,* lacks the genomic signature of the specialized genome defense mechanism called Repeat Induced Point mutations (RIP). We identified the centromeric regions in *P. anserina* and *P. pseudocomata* using chromatin immunoprecipitation targeting the centromere-specific histone variant cenH3, and using comparative genomics we inferred the size of centromeric regions in the other species. We found that while the centromere structure in the complex is generally well conserved, the centromeric regions of *P. pseudocomata* has gone through a rapid change. Specifically, the size of the centromeres in *P. pseudocomata* are 35-46 kb, which is significantly smaller than those of the other species (44-90 kb), and the DNA-transposon *discoglosse* is the most abundant TE family instead of the typical LTR-retrotransposon *crapaud*. Taken together, our data strongly indicates a link between genome defense and centromere evolution in fungi.

## Introduction

Transposable elements (TEs) are mobile elements that proliferate by making new copies of themselves within the genome (1). Throughout eukaryote evolution, many genomic host-defense systems have evolved to counteract the negative effects of these (2–4). Within the kingdom Fungi, multiple specialized defense mechanisms have evolved (3). One of these is the repeat induced point mutations (RIP)-mechanism that induces C-to-T mutations into any repeated region in the genome (5–8, 3, 9, 10). RIP is widespread in the phylum Ascomycota, and acts as a key mechanism for maintaining small and streamlined fungal genomes by limiting TE accumulation. At the same time, TEs often play an important role for genome evolution in fungi. For example, in many species they build up important genomic regions such as centromeres (11). Centromeres are regions on chromosomes where the kinetochore binds to assemble microtubules during cell division (12). They are ubiquitous among eukaryotes and essential for functional chromosome segregation. Notably, the DNA of the centromeres is highly variable among eukaryotes, and the contrast between the conserved function and rapidly evolving DNA sequence and components has been referred to as “the centromere paradox” (13). Our understanding of centromere sequence composition has recently improved thanks to major advances in both sequencing and assembly technologies. Studies in animals (14–16), plants (17–19), and fungi (20, 21, 11) have contributed to a deeper understanding of both the functional role of centromeres in cell division and their role in genome evolution. From the fungal species for which centromeres have been characterized, it is clear that fungal centromeres show high variability in both size and content: they range from small point centromeres of ~125 bp found in the *Saccharomyces* species, to regional centromeres that can be several hundred thousand base pairs long and often filled with TEs (11). Because many fungal centromeres are composed of TEs, the evolution of a wide-array of specialized genome defense mechanisms against TEs (3) should impact centromere evolution. Hence, the interaction between TEs and genome defense needs to be integrated into the study of fungal centromeres (21).

In this study, we use the *Podospora anserina* species complex to investigate the link between genome defense, centromere size, and a shift in transposable element composition. This species complex is made up of eight recently diverged species belonging to the Ascomycete fungal order Sordariales (22, 23). The complex is most known from *P. anserina*, which has been used as a model species in the study of various genetic and cell biology phenomena (24), including the molecular underpinnings of RIP (25–27). The recent release of high-quality long-read assemblies of the genomes of seven species in the *P. anserina* complex (28–30) together with a solid background understanding of their TE composition (28, 31–33), offers an ideal opportunity to study centromere evolution over short evolutionary timescales, and in particular the signature of interaction between repetitive elements and genome defense in this region.

First, by investigating the relationship between patterns of RIP and repeats in the genomes, we found a clear lack of typical signature of RIP in one of the species, *P. pseudocomata*, suggesting the loss of the defense mechanism in this lineage. Furthermore, we defined the centromere region in *P. anserina* and *P. pseudocomat*a by chromatin immunoprecipitation targeting the centromere-specific histone variant cenH3, and using comparative genomics we inferred the size and location of the centromeric regions in the other species in the complex. With a comparative genomic approach, we found *P. pseudocomata* to have a drastic shift in both size and content of its centromeric regions that has taken place within the last 1 MY, suggesting a strong connection between RIP and centromere structure.

## Results

### The *Podospora pseudocomata* genome lacks the typical signature of RIP

Given that RIP hypermutates repetitive DNA with C-to-T mutations, a negative association between GC-content and repeat proportion in species with an active RIP machinery is expected. We analyzed the association between the proportion of GC-content and repeat proportion in sliding, non-overlapping windows across the seven genomes of the *P. anserina* species complex, and for all species except *P. pseudocomata,* we found a strong negative association (Linear regression model, P <0.001 for all species, Adjusted R^2^ value for *P. pseudocomata* = 0.05 and for the others 0.51-0.71) (Figure 1A). For reference, the annotated repetitive content of the seven genomes in the species complex range between 3.76 and 7.1%, and for this trait, *P. pseudocomata* (with a 4.82 % repeat content) does not deviate from the other species (Supplementary table 1). By using a RIP-index analysis, which is based on calculating ratios of RIP-related dinucleotide frequencies, we show that 0.15% of the genome of *P. pseudocomata* had been affected by RIP, while the corresponding percentage in the other species’ genomes ranged from 1.92 % to 6 % (4.76 % in *P. anserina*, where RIP is experimentally known to be active). This suggests that RIP is active in all species except in *P. pseudocomata*, and since RIP is thought to be ancestral to ascomycetes (34, 35) our assumption is that it has been lost in the *P. pseudocomata* lineage.

**Figure 1:**
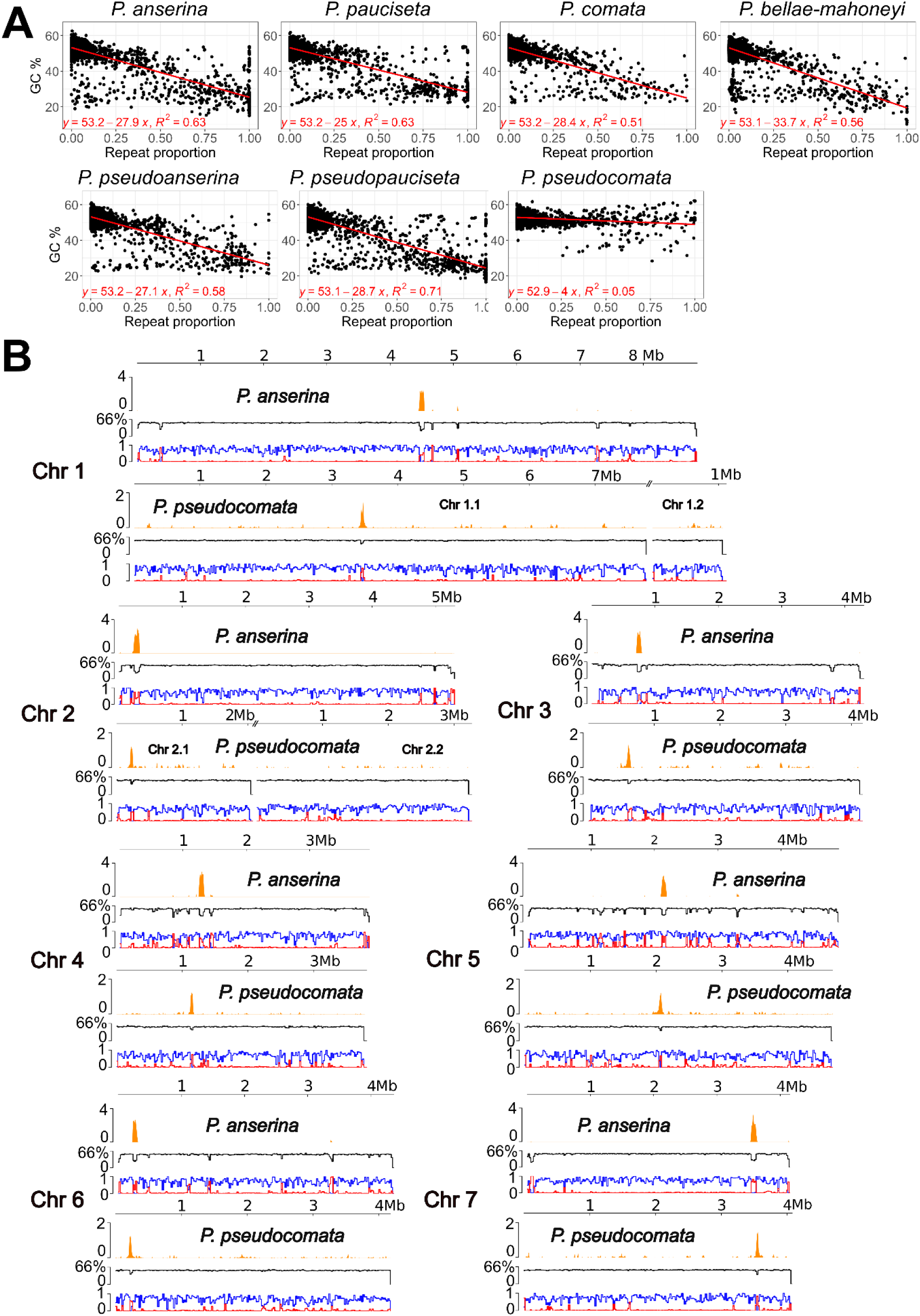
Lack of RIP signal in *Podospora pseudocomata* and localization of centromeric regions in *Podospora anserina* and *Podospora pseudocomata*. A) Comparison of GC-content and repetitive content in 5kb non-overlapping windows. B) Localization of cenH3 in *P. anserina* and *P.pseudocomata*. Tracks from top to bottom: 1) Enrichment of immunoprecipitated cenH3 compared to input (*P. anserina*) or mock (*P. pseudocomata*). 2) GC % calculated in 20kb windows 3) Gene (blue) and repeat (red) proportions calculated in 20kb windows.

The primary pathway of RIP is thought to act through a cytosine methyltransferase encoded by the gene *rid* (36). We located an intact ortholog of *rid* (QC762_119440) in *P. pseudocomata*, which differs by eighteen variable sites (eight of which are non-synonomous) from the ortholog in *P. anserina,* (Pa_1_19440) (27). Four of the non-synonomous changes affect the first half of the CDS, and the last four affect the 3’ end. All eight non-synonomous changes have no impact on the conserved domains essential for the enzymatic activity of RID. (data not shown). Hence, we did not reveal insights into the mechanism behind the loss of RIP by comparing the gene sequences between species.

We further found that the lack of RIP is a trait inherent to the species *P. pseudocomata* and not a strain-specific trait of the reference strain. We approached this topic by sequencing and assembling a newly discovered second strain of *P. pseudocomata*, IMI230595m, sampled from another continent. We found that 0.23% of the genome is RIP affected in this strain and that it similarly has an intact ortholog of *rid*. No other *P. pseudocomata* strains are known, but the consistent low proportion of the genome that is affected by RIP in the two known strains strongly suggests that the pattern we observe is a species-specific trait (Supplementary Table 1).

### Identification and first characterization of the centromeric regions in the *Podospora* genomes

Despite its extensive use in biological research and long use as a model system, the centromere location in *P. anserina* has so far only been inferred from genetic maps (24, 37), and the centromere size and location has until now been completely unknown in the other species of the complex. In fungi, like in other eukaryotes, centromeres are typically characterized by the presence of a histone H3 variant, cenH3, also known as CENP-A (11). In order to be able to investigate the interaction between repetitive elements and genome defense in the centromeric region of *Podospora*, we first choose to precisely localize the centromeric regions of *P. anserina* and *P. pseudocomata* by identifying the centromere specific histone variant cenH3 (Pa_7_8690 in *P. anserina*). The protein was tagged with GFP and expressed into the two species. We then inferred the genomic localization of cenH3 using chromatin immunoprecipitation of the GFP tag. Through sequencing (ChIP-seq), we were able to identify significantly enriched peaks for both *P. anserina* and *P. pseudocomata* (Figure 1B, Supplementary Figure 1). In both species, there was one clear enriched peak for each of the seven chromosomes corresponding to large repetitive element islands and a depletion of GC-content (Figure 1B). In addition, in *P. anserina*, the cenH3 peaks were found at the expected locations given previous linkage maps (32).

We defined the centromeric region borders at the genes flanking the repeat islands, which for most coincided with a drop-off of cenH3 enrichment. The exception to this pattern was CEN2 in *P. pseudocomata* where there was no cenH3 enrichment at the repeats at the downstream end of the centromeric region (Supplementary figure 1). With lower thresholds for the mapped reads the enrichment at the right flank was present and strong (data not shown), i.e., mapped reads at this downstream location had been removed through read mapping filtering. Based on that result, we opted to keep the CEN2 border at the gene flank for further analyses. We then used the flanking genes in *P. anserina* and *P. pseudocomata* to locate the orthologs in the other species. Our data suggest that the placements of the centromeric region on the chromosomes are conserved in the species complex. Specifically, we identified AT-rich TE islands with flanks that are syntenic throughout the species complex with only minor rearrangements of genes. Hence, for further analyses, we assumed that the regions between these flanking genes correspond to the centromeric regions throughout the species complex.

### Multiple factors behind a reduced GC of the centromeric regions

When contrasting the patterns of GC-content and repeat proportion between centromeric and non-centromeric regions, three main patterns were identified: First, all species showed a lower GC-content in the centromeric regions than in the rest of the genome (Supplementary Figure 2), a commonly observed feature of eukaryote centromeres (12). Second, *P. pseudocomata* had an overall higher GC-content in centromeric regions (41%) than the other species, which all showed a range between 27-28%. In accordance, RIP-index analysis showed that the proportion of RIP inside the combined centromeric regions was 15.8 % for *P. pseudocomata* and ranged between 64.4-76.3 % for the other six species. Third, the pattern of the centromeric region contrasted the rest of the genome in that we did not observe a negative linear relationship between GC and repetitive proportion in any of the species (Supplementary Figure 2). Noteworthy, in *P. anserina* we even found a slight positive association (Linear regression model, p = < 0.01, R^2^ = 0.23; Supplementary Figure 2). Taken in combination, these patterns reveal signatures of RIP in the centromeric regions, but also indicate that there is an additional factor other than RIP driving down the GC-content inside the centromeric regions of all species. Note that it is difficult to disentangle whether the weak signal of RIP we observed in *P. pseudocomata* is due to RIP still being active at low levels or if there are old copies of TEs remaining that carry RIP signature, *i.e.,* it was inherited from before the loss of RIP.

### Comparative analyses suggest a rapid evolution of centromere size and content in *Podospora pseudocomata*

Visual inspection of the aligned centromeric regions (Figure 2A) revealed that the synteny of the centromeric regions reflected the phylogeny of the species complex, with the two closely related sister species *P. pseudoanserina* and *P. pseudopauciseta* showing the highest synteny, followed by the sister species *P. anserina* and *P. pauciseta*, while each pairwise comparison representing a more ancient split showed little or no synteny. In addition, the centromere flanking gene regions were conserved in all chromosomes except CEN3 in *P. pseudocomata*, where two copies of the LTR retrotransposon *rana* had inserted at the right flank (Figure 2A), but outside the cenH3 enrichment peak of that centromeric region (Supplementary figure 1). Overall, these results show that the evolution of the centromeric region in the *P. anserina* species complex is tractable over the investigated evolutionary timescale.

**Figure 2:**
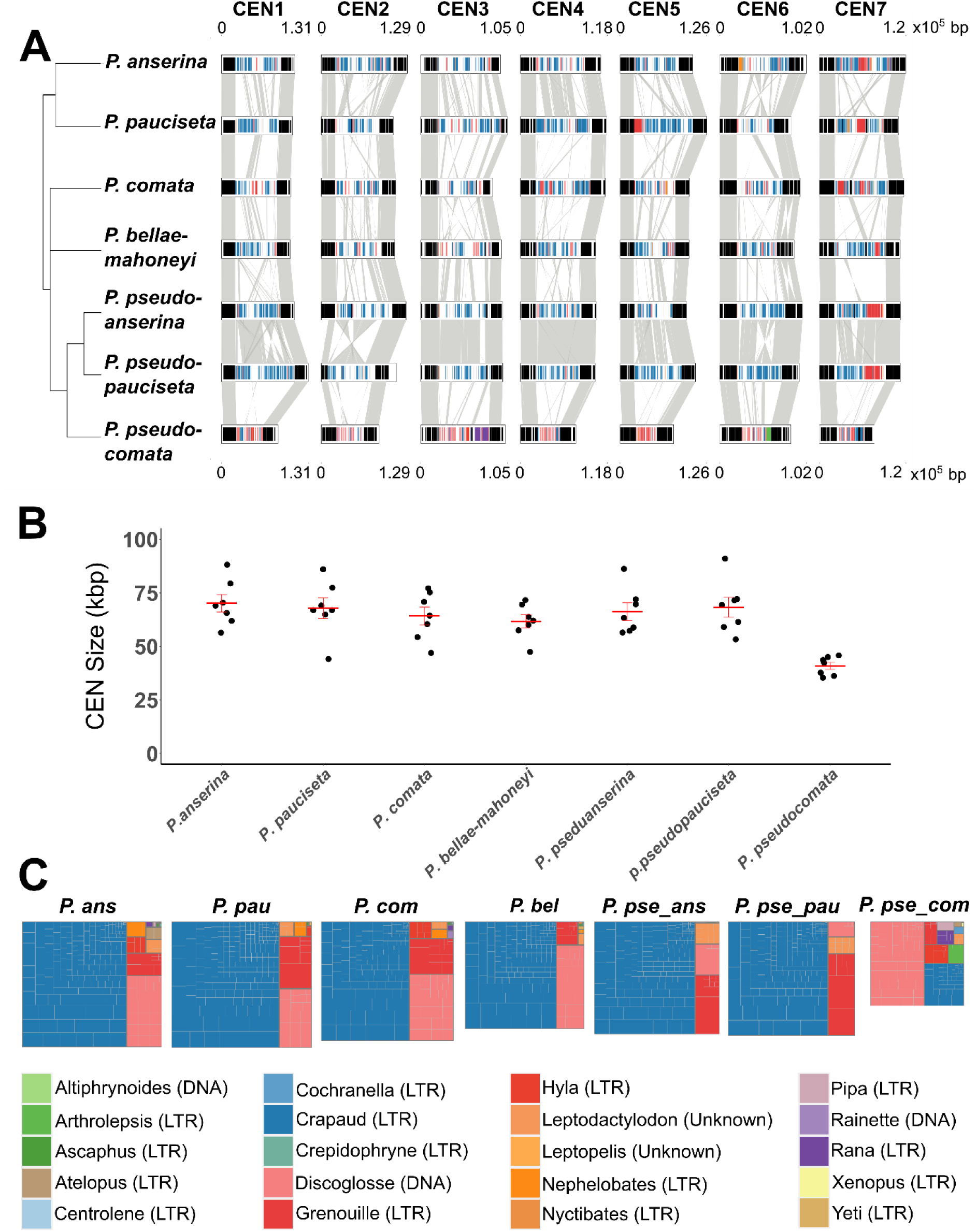
Comparison of the centromeric regions in the *Podospora anserina* species complex. A) Synteny of the centromeres plus 20 kb flanks, with transposable elements highlighted in color (see legend at the bottom of figure) and genes shown in black. Gray links indicate Nucmer alignments. The cladogram is a modified version from Ament-Velásquez *et al* (2024); as no suitable outgroup is available, the tree is rooted with *P. anserina* and *P. pauciseta* B) Jitter plots of centromeric region sizes of the seven chromosomes of each strain. Standard error bars and means are shown in red C) Treemap tiles of transposable element families inside the centromeric regions of respective species. The classified Order of the TE are shown in parentheses. Overall size of the treemaps have been scaled to represent the overall transposable element amount in the centromeric regions. P. ans = *P. anserina*, P. pau = *P. pauciseta*, P. com = *P. comata*, P. bel = *P. bellae-mahoneyi*, P. pse_ans *= P. pseudoanserina*, P. pse_pau *= P. pseudopauciseta*, P. pse_com *= P. pseudocomata*.

Investigating the size of the centromeric regions revealed that the size of the centromeric regions of the species in the complex are well conserved, with the exception of *P. pseudocomata*, where the centromeres are significantly smaller than those of the other species (Figure 2A and B). The median size of the centromeric region in *P. pseudocomata* was found to be 42 kb, while the medians of the other species’ range between 62-69 kb (Kruskal-Wallis chi-squared test, χ^2^ = 18.691, p = 0.0047). Furthermore, we found that there was little intraspecific variation in centromere sizes of both the two *P. pseudocomata* strains and between *P. anserina* strains (Supplementary Figure 3, Supplementary Table 2), which suggests that the difference in centromere sizes are not a strain-, but rather a species-specific trait. Furthermore, both the overall TE content and the centromeric TE content are similar in the two investigated genomes of *P. pseudocomata*, with the main difference being a non-centromeric expansion of the LINE family *kermit* in the reference strain CBS415.72m which was not found in IMI230595m (Supplementary figure 4), strongly suggesting that they are two genetically diverged strains of *P. pseudocomata* and not clones. Specific analysis of the TE content of the centromeric regions showed that *P. pseudocomata* had both a lower total number of TEs and a different TE composition than the other species. Specifically, a DNA-transposon family, *discoglosse*, was the most abundant TE family instead of the otherwise most abundant LTR-retrotransposon *crapaud* (Figure 2C). The absence of a suitable outgroup for the species complex makes it difficult to infer directionality of evolutionary events, but we assume that the difference in size is a result of a shrinkage in *P. pseudocomata’*s centromeric regions rather than an increase in size in the other species of the complex, and that a turnover of the TE composition has taken place in the *P. pseudocomata* lineage.

The shift in centromere architecture is likely to have occurred recently in evolutionary time, given the close relationship of the species in the complex (as evidenced by a high genome collinearity, low overall divergence levels (>98% identity in genic regions; (30)), and incomplete reproductive isolation (38, 39)). In addition, the short internal branches separating the *Podospora* species in the phylogeny suggest a rapid, nearly simultaneous diversification of all lineages (30). Hence, we opted for estimating the divergence time of the whole species complex as an upper bound limit to the age of the centromere changes in *P. pseudocomata*. We used published estimates of synonymous substitution rates (*d_S_*) between *P. anserina* and all other lineages in 100 random genes (39) as a proxy of neutral evolution. Assuming a simple mutation-drift equilibrium model, 24 generations per year, and two different fungal neutral substitution rates (40) as minimum and maximum, we estimated that the species complex diverged less than 1 Mya (between 38 469 to 713 808 years ago using the different substitution rates (Supplementary table 3). Despite the limitations of this method, the upper bound of the obtained range highlights the recency of *P. pseudocomata*’s genomic shift. As a comparison, a similar case of changes in centromere size and composition, along with the loss of a genome defense mechanism can be seen between *Cryptococcus* species (21), which diverged 34 Mya (41).

### Depletions and enrichment of TE families inside the centromeric regions

We further investigated the distribution of different TE families in the genomes of the *P. anserina* species complex, and found that the number of TE families present in the centromeric regions is lower than in the rest of the genome for all seven species (Figure 3A, Supplementary tables 4-10). Permutation tests of the centromeric windows versus other repeat-rich windows for each genome revealed that certain elements were found to be significantly depleted in the centromeric regions of the different species. None were depleted in all species but, for example, both the *hyla* (Copia LTR) and *grenouille* (Ty3 LTR) repeats were depleted in the centromeric regions of *P. anserina* and *P. pseudopauciseta* (supplementary table 5 & 10), and two other TE families were found to be depleted in the centromeric regions of one species each: *centrolene* (Copia LTR), in *P. pauciseta*, and *pelobate* (Tc1/Mariner DNA transposon), in *P. comata* (Supplementary table 6). Furthermore, there were four TE families for which copies were enriched in the centromeric regions (Supplementary table 4-10). Of these, the *crapaud* family was enriched in the centromeric regions of all seven species; *discoglosse* was enriched in three species; *grenouille* in two species; and *hyla* in one species (Supplementary tables 4-10). *P. pseudocomata* was the only species in which all of these four TE families were enriched in the centromeric regions (Supplementary table 4).

**Figure 3:**
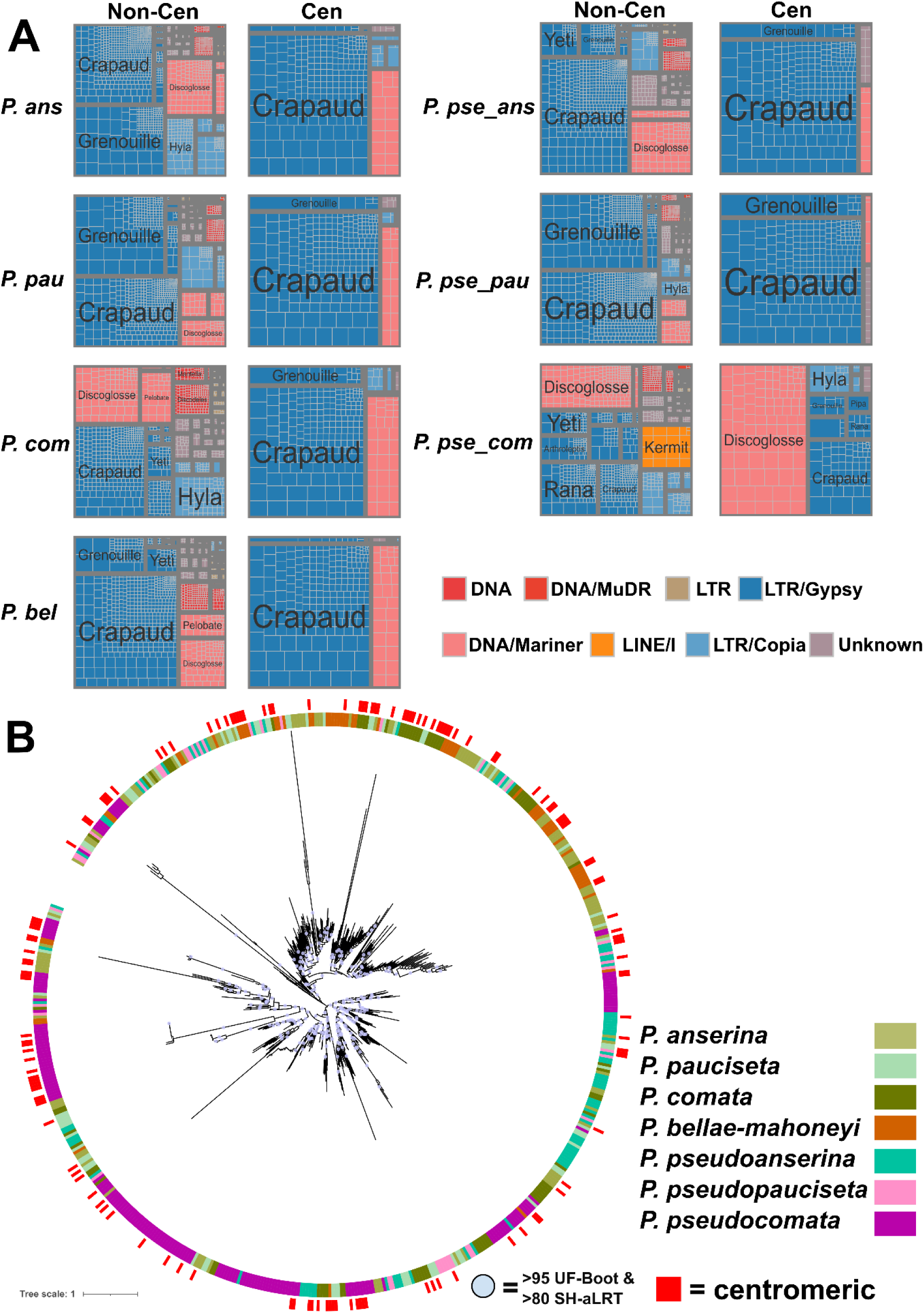
Distribution of TEs inside and outside the centromeric regions and expansions of TEs. A) Comparison of proportions of TE-families present at centromeres (Cen) and outside of centromeres (Non-Cen) in the different Podospora genomes. B) Maximum likelihood tree of the copies of the discoglosse DNA transposon in the species of the species complex (inner ring), and centromeric copies (outer ring). P. ans = *P. anserina*, P. pau = *P. pauciseta*, P. com = *P. comata*, P. bel = *P. bellae-mahoneyi*, P. pse_ans = *P. pseudoanserina*, P. pse_pau = *P. pseudopauciseta*, P. pse_com = *P. pseudocomata*.

We recently classified the *crapaud* family into 14 different subfamilies based on the LTRs (33). We next sought to investigate whether there were differences between these subfamilies in terms of centromeric regions depletion and enrichments. None of the subfamilies were depleted in centromeric regions compared to other repeat rich regions in any of the genomes. However, we found that there were differences in enrichment between subfamilies. Some subfamilies (LTR2, LTR4, LTR9, LTR10, and LTR14) were enriched in the centromeric regions of more than five species while others (LTR1, LTR3, LTR5, LTR6, LTR7, LTR8) were enriched in two or fewer species (Supplementary table 11). A maximum likelihood phylogeny of the subfamilies support that all subfamilies have both centromeric and non-centromeric copies (Supplementary figure 5).

### TE dynamics in *Podospora* reveals past RIP patterns and multiple recent TE expansions in *P. pseudocomata*

The lack of centromere specific clusters in *crapaud* phylogeny (Supplementary figure 5) supports our previous finding that the *crapaud* subfamilies mainly diversified in the ancestor of the species complex (33), and that more recent, genomewide, *i.e*., non-centromere-specific, smaller bursts of the subfamilies have taken place within species (Supplementary Figure 5). We similarly wanted to investigate *discoglosse* as it is the most abundant TE in the *P. pseudocomata* centromeric regions. A maximum likelihood phylogeny of *discoglosse* showed large clusters of TE copies originating from the *P. pseudocomata* genome, indicative of multiple expansions in this species after the split from the other species (Figure 3B). Some of these expansions are very recent, as suggested by clusters with very short branch lengths in the phylogeny, while others consist of more diverged copies. All of these clusters have copies both inside and outside the centromeric regions. This contrasts with what we observe for *crapaud* and indicates that *discoglosse* has had relatively recent and more substantial transpositions in *P. pseudocomata*, including into centromeric regions. Yet, bursts of *discoglosse* in *P. pseudocomata* were, similar to *crapaud*, not confined to the centromeric regions.

To continue our TE investigations, we turned to examining individual families of TEs in *P. pseudocomata*. Despite the overall RIP index values being low in the genome, we found that some TE families in *P. pseudocomata* have a higher RIP composite index than others (Supplementary figure 6A), ranging from 1.6 in the single solo-copy of *hyla* in the genome (we considered composite index values over zero to indicate RIPped copies) to −1.395 in *Rainette* (indicative of no RIP). Given the non-discriminatory activity of RIP, this is most likely explained by past TE family expansions before the loss of RIP and that the copies have remained in the genome. As expected from the mutational bias introduced by RIP, the impact of RIP mutations was also reflected in the distribution of GC-content of the TE families (Supplementary figure 6B). In addition, we see that copies of certain families show a wide range of RIP and GC variation, for example both *crapaud* and *grenouille*. However, many TE families had average RIP composite index values below zero (Supplementary figure 6A), and limited GC-content ranges (Supplementary figure 6B).

Since RIP levels are overall low in *P. pseudocomata*, we calculated average copy sequence divergences (Kimura2) from the representative sequences of all TE families in *P. pseudocomata*. This type of analysis takes the transition/transversion ratio between the consensus and copies into account to determine the age of these copies. Since RIP skews these ratios, this analysis is typically difficult to perform in species that have RIP, which is why it has not been done before for the *P. anserina* species complex. *Discoglosse* stood out as it had a high genomic proportion in *P. pseudocomata*, and intermediate divergence levels (Supplementary Figure 6C), meaning that it likely proliferated both before and after the loss of RIP. C*rapaud* instead showed high average divergence (Supplementary figure 6C) and hence likely represents primarily ancient proliferation, prior to loss of RIP. We also found that the *kermit*, *rana*, *arthroleptis*, *sasquatch*, and *xenopus* families all had lower average Kimura2 divergences than *discoglosse* and also relatively high genomic proportions (Supplementary figure 6C), suggestive of very recent expansions of these families in *P. pseudocomata*, post loss of RIP. Furthermore, maximum likelihood phylogenies of the LTRs of the *rana, arthroleptis*, and *xenopus* LTRs suggest that the expansions of these three families are primarily found in in *P. pseudocomata* (Supplementary figures 7-9) and *kermit* and *sasquatch* families lacked any copies in any of the other species. Taken together, there are specific TE expansions of certain TE families throughout the genome of *P. pseudocomata*.

## Discussion

Centromeres are ubiquitous features of eukaryotes, and understanding their evolution is crucial for understanding forces driving genome evolution. The *P. anserina* species complex offers an attractive opportunity to investigate short term evolution of centromeres, as it consists of multiple, closely related species that diverged from each other less than 1 Mya, and for which high-quality genomic information is available. In particular, the finding of loss of RIP in *P. pseudocomata* made it possible for us to infer the association between genome defense and centromere evolution over this short evolutionary timescale. Even if we, with this data, are not able to directly infer causality, we report a strong correlation between loss of RIP and a drastic reduction in centromere lengths and a high turnover of transposable elements in the centromeric region of *P. pseudocomata*.

RIP is widespread in ascomycete fungi, however, previous large-scale studies have shown that it may occasionally be lost (42, 43). We found that *P. pseudocomata* shows no indication of active RIP genome defense mechanism. This is one of few detailed reports of a loss of RIP and the consequences in a species. As the *rid* gene appears functional, the exact mechanism of RIP loss is still unknown, and needs further investigation. In terms of consequences, one could imagine a scenario of loss of defense against TEs resulting in an increase in TEs. For example, in a recent study, the loss of RIP in the class *Leotiomycetes* was suggested to be related to a genome size expansion due to TE proliferation (43). In contrast, we instead conclude that *P. pseudocomata* may have higher diversity of active TE families, which could potentially be driven by loss of RIP. However, the TE load in terms of abundance is among the lowest in the species complex (30, 33). The low TE abundance is similar to what is found in *Sordaria macrospora*, a species that also lacks RIP, and in which the genome has lower TE load than its close relative *Neurospora crassa* (44–46). In *Schizosaccharomyces pombe* the loss of RNAi, a different type of defense mechanism against TEs, triggered LTR retrotransposon repression by CENP-B homologues (47). It is similarly possible that the loss of RIP in *P. pseudocomata* has triggered the activation of another TE repressing system, like RNAi or CENP-B associated inhibition. Another hypothesis is that TE load in *P. pseudocomata* is instead reduced due to other processes, for example demographic processes and/or strong purifying selection, which would relax the selective pressure to keep up a costly TE defense and allow it to be subsequently lost. Yet another hypothesis is that RIP is lost as a result of selection for increased evolutionary rate, which was speculated to have been driving the loss of RNAi in the basidiomycete human-pathogenic fungus *Cryptococcus deuterogattii*, to overcome host immune defense (21). None of the species in the *P. anserina* species complex have been observed to be pathogenic, but are assumed to primarily be coprophilous, meaning that they need to grow on the dung of herbivores as part of their life cycle. This environment is very competitive, and may lead to a lifestyle that is stressful in similar ways as a pathogenic lifestyle and thus present similar advantages to losing genome defense in order to increase the evolutionary rate of the genome. However, members of the species complex have been isolated from other environments, such as soil and as an endophyte, suggesting that dung is the preferred but not unique biotype (22, 48). The *P. pseudocomata* reference strain, CBS415.72m, was in fact isolated from soil. Apart from *P. anserina*, the ecology of the species complex is poorly studied. One could imagine that *P. pseudomata* has a life-style or faces some other ecological challenge than its closely-related lineages, and that this is connected to the loss of RIP. To be able to possibly link lifestyle with the genomic patterns we have observed, more studies are needed on the biology of the rest of the species complex.

Remarkably, the centromeric regions of *P. pseudocomata* are almost half the size of the rest of the *P. anserina* species complex. They also have a higher TE density, and have had a turnover of centromere dominant TE family, to the DNA transposon *discoglosse* instead of the LTR retrotransposon *crapaud*. The changes in *P. pseudocomata* emphasize that there are no specific requirements for either size or content for maintaining centromere function. The co-occurrence of loss of genome defense and shrinkage of the centromere in *P. pseudocomata* is similar to what is reported from *C. deuterogattii* that similarly displays smaller centromeres and also truncated TEs (21). Several recent studies on centromeres have described centromere retrotransposon enrichment and observed TE-related transcriptional activity within centromeres, although the exact nature of this activity remains unclear (49–51). Investigating transcriptional activity inside the centromere regions of *P. anserina* and *P. pseudocomata* could thus be a future avenue to investigate the function of centromere transcription and the impact of a shift in centromere TEs.

### Centromeric TE turnover and the interaction between TEs and centromeric regions

From our results we see a fast turnover of TEs inside the centromeric regions of *P. pseudocomata,* from *crapaud* to *discoglosse.* In addition, we observe that the TEs that are enriched inside the centromeric regions are a distinct subset of those in the rest of the genome, with some TE families never present in centromeric regions. Centromere associated TEs have been found in both fungi, animals, and plants (52–56). Several of the non-centromeric families have also had recent transpositions in *P. pseudocomata* meaning that it is not only the families with the latest expansions that are found in the centromeric regions. Non-random distribution of TEs have previously been described as a balance between integration site preference of the TE and selection against the inserted TE by the host (57). In compact genomes, such as many fungi, integration is often limited to heterochromatic regions or into tDNA or rDNA (57). Examples of preferential insertion sites are the Ty3 families, which carry a chromodomain with heterochromatic DNA as preferred insertion sites (58–60) or the P-element, a DNA-transposon, in *Drosophila melanogaster* that preferably inserts at replication origins (61). In the species complex, we observed chromodomain-carrying Ty3 families that were either enriched inside centromeric regions (like *crapaud* and *grenouille*), or not (like *yeti* and *xenopus*). Another explanation for this difference could then be a difference in purifying selection between TE families inside or outside of centromeric regions leading to the bias we observe. As mentioned above, AT-rich DNA is favored at eukaryotic centromeres (12), which means that new TE-insertions with higher GC may be subject to purifying selection. For the *crapaud* family, we previously reported 14 different subfamilies based on the LTRs (33), here we see a difference in enrichment inside the centromeric regions between them. However, there are no species-specific centromeric region clusters of copies in any of the subfamilies. One of the main findings of our previous study of *crapaud* was that the diversification of the subfamilies predated the species complex diversification (33). The lack of species-specific expansions in the centromeric regions could thus be explained by *crapaud* invading the centromeric regions mainly in the ancestor of the species complex. It is therefore still an intriguing possibility that the subfamilies have evolved to have differential insertion patterns. The varying U3 region at the LTRs may carry some important transcription factors or other regulatory elements (33, 62), which may alter the insertion preference.

## Conclusion

In this study we have successfully localised the centromeric regions using ChIP-Seq in two species of the *P. anserina* species complex and inferred the positions in the rest of the species complex through preserved synteny. We have described the loss of genome defense and the impacts it has had on the dynamics of TEs and centromeric regions in *P. pseudocomata*. It has previously been shown that centromeric DNA is affected by changes in RNAi-based genome defense. Here we show that this link between centromeric regions and genome defense likely extends further to encompass different types of TE defense mechanisms. We observe a fast turnover of TEs in the centromeric regions of *P. pseudocomata* after loss of RIP, which indicates that understanding the interaction between TEs and genome defense is important for studying the evolution of centromeres in fungi, and possibly other eukaryotes as well. In addition, studying centromere evolution in the *P. anserina* species complex, where the species have short divergence times, highlights the rapid evolution of centromeres in eukaryotes and that studying closely related organisms is needed in order to observe the diversity of centromere structures and the extents of centromeric turnover events.

## Materials and Methods

### Strains and media

Full information of all strains used in the genomic analyses can be found in supplementary table 1. The *P. anserina* strain used in the wet lab experiments in this study is the laboratory S strain (63, 32), for which transformation methods and laboratory protocols are well developed. For the bioinformatic analyses we instead used the Wa63^+^ strain (64), which has the more contiguous assembly PaWa63p (28), and shares a high level of sequence conservation with the S strain. The *P. pseudocomata* strain CBS415.72 was used in both the wet lab and the bioinformatic parts of the study (30). We generated and assembled a genome of *P. pseudocomata* (strain, IMI230595, see details below) specifically for this study. This strain was originally isolated by R.F. Cain, D. Griffin & JC Krug from *Bos taurus* dung located in Kenya in 1978. We ordered this strain from CABI where it is listed as *P. pauciseta*, but after re-examining the strain we identified it as *P. pseudocomata.* Media composition used during the wet lab portion of the experiment can be found in Silar 2020 (22).

### Culturing, sequencing, and assembly of *P. pseudocomata* strain IMI230595

High molecular weight DNA was obtained from *P. pseudocomata* IMI230595 according to the protocol outlined in (65). Specifically, strains were grown in 3% liquid malt extract for 5 days. Mycelia were filtered from solution and freeze dried overnight. A sterile mortar and pestle was used to grind 1g of dried mycelia. A Genomic TipG-500 column (Qiagen) was used to extract DNA following the manufacturer’s protocol. The extracted DNA was sequenced using the PacBio Sequel II Technology Platform using the multiplexing protocol on 1 SMRT Cell 8M at ScilifeLab, Uppsala, Sweden.

The CCS PacBio reads were assembled using hifiasm v. 0.19.5 (66). The resulting scaffolds were oriented to match the chromosomes of the reference genome of the strain S (32) by using MUMmer v. 3.23 (67) alignments. Small scaffolds corresponding to ribosomal DNA and mitochondrial fragments (mainly α senDNA) were discarded. The final assembly contains the seven chromosomes in a single scaffold, some of which have telomeric repeats on one arm (chromosomes 1, 4, and 6) or both arms (2, 5, and 7). Annotation of protein coding genes was performed as in Ament-Velásquez et al. (2024). Briefly, *de novo* gene models were produced with MAKER v. 3.01.04 (68, 69). The PODANS_v2016 annotation (70) and the *P. comata* reference annotation (71) were used as external protein evidence, while external transcript models were obtained from published RNA-seq data of *P. anserina* and *P. comata* (28, 29). Functional annotation was performed with Funannotate v. 1.8.15 (72).

### Chromatin immunoprecipitation in *P. anserina* and *P. pseudocomata* and ChIP-sequencing

To verify the positions of the centromeric regions in *P. anserina* (strain S) and *P. pseudocomata* (strain CBS415.72), we transformed the strains with a plasmid carrying the cenH3 histone gene fused to the eGFP CDS to express a C-terminally GFP-tagged cenH3 histone. For each strain, a synthetic construct was ordered composed of the cenH3 coding sequence without stop codon followed, in frame, with three glycine codons (ggc, as linker) and the GFP coding sequence (Thermofisher Scientific Invitrogen GeneArt gene synthesis service).

For *P. pseudocomata* CBS415.72, the construct was PCR amplified using primers containing linker sequences that carried *Spe*I and *Xho*I restriction sites (See supplementary table 12 for primers). The PCR product was digested with *Spe*I and *Xho*I enzymes and ligated into a plasmid carrying resistance to hygromycin (pBC-Hyg) (73) linearized with the same enzymes. For *P. anserina* strain S, the construct was PCR amplified with primers containing *Bam*HI and *Eco*RI restriction sites. The PCR product was digested with *Bam*HI and *Eco*RI and ligated into the pAKS120 plasmid linearized with the same enzymes (27). All constructs were checked by sequencing. The plasmids were transformed into their respective species in wild-type strain as described in (22, 74). The transformants were all checked by PCR for the presence of the GFP coding sequence. The ChIP experiment was performed as in (75). One transformed strain of each species plus wildtypes were selected and grown out in liquid M2 medium at 27℃ for 2 days in roux flasks. For *P. anserina*, we also included an extra technical replicate for the transformed strain. The chromatin immunoprecipitation was then performed following the protocol in (74).

The chromatin of two pooled ChIP experiments of *P. pseudocomata* and the chromatin of the mock strain without cenH3-GFP were sent to the SciLife-lab sequencing facility for library preparation and sequencing. The sequencing was done using the SMARTer ThruPLEX DNA-seq, which includes library preparation for small concentrations of DNA and sequencing using 300x paired-end Illumina reads. The chromatin of two *P. anserina* ChIP experiments, input, and mock, respectively, was sent to Novogene ChIP-seq service at a later date, which included library preparation and sequencing using Illumina NovaSeq X plus platform using paired-end 150bp reads.

### Bioinformatic processing of ChIP-seq data and analysis of chromosomal contents

The sequenced Illumina reads were first preprocessed with FastP v 0.23.4 (76) to remove duplicate reads and then trimmed using Trimmomatic v0.39 (77) with the settings: [ILLUMINACLIP:data/TruSeqadapters.fa:2:30:10:8:true SLIDINGWINDOW:4:15 LEADING:30 TRAILING:30 MINLEN:50]. The reads were then mapped to the *P. anserina* long-read assembly strain PaWa63p (28) and the *P. pseudocomata* assembly of the strain CBS415.72m (30). PaWa63p is a high-quality assembly without any tracks of Ns unlike the reference assembly of the S^+^ strain. To map the reads we used BWA-MEM v.0.7.17 (78) and then Samtools v1.2 (79) was used to save the mapped reads to BAM-files. Then reads were filtered using the command [Samtools view -F 256 -q 30 -e ’[AS]>=145’]. The bamCompare command from deepTools v3.3.5 (80) was used to compare the mapped ChIP-seq reads of the cenH3-tagged samples with the controls (input for *P. anserina* and mock for *P. pseudocomata*) using the commands [--binSize 20 --normalizeUsing BPM --smoothLength 60 --extendReads 150 --centerReads]. Peak calling was performed using MACS2 v2.2.9.1 (81) using q=0.01 and the respective genome sizes for the two assemblies. Masked repeat content was obtained using RepeatMasker v 4.1.5 (82) using the latest updated repeat library (33). GC-content, gene-content and repeat-content was calculated in 20 kb sliding windows using a custom script (https://github.com/Ivwster/Podospora_Centromeres). The bigwig files, gene-content, GC-content, and repeat-content were visualized using Pygenometracks v3.8 (83). For visualization we picked the cenH3 ChIP IP from one of the two experiments, but the pattern was the same in both (data not shown). To estimate levels of RIP in the genomes we used the default settings of web tool TheRIPper (84), and used the “RIP affected genomic proportion” estimation of the RIP profile provided by the software.

### Bioinformatic retrieval and alignments of the centromeric region proxies in the species complex

To bioinformatically collect the flanking genes and positions of the other species from the position in *P. pseudocomata* centromeric regions, we first located the orthologous genes flanking these regions in the other species using BLASTn, annotation data, and IGV v2.14.1 (85) (Supplementary table 2). We then extended the flanks 20 kb on each side, with the exception of *P. pseudocomata* centromere 3 where we extended 40 kb on the right flank due to TE-insertions. We then pairwise aligned each pair of species according to the phylogeny (30) using the NUCmer algorithm from MUMmer v3.23 (86) with the parameters: [--mum -b 400 -g 150 -o -l 15] and filtered for percent identity above 0.8. For each chromosome in each species, we then verified the presence of a repetitive island, which all were among the largest repetitive islands in each chromosome. The centromeric regions and alignments were visualized in R v4.3.2 (87) using the ggplot2 v3.4.4 (88) and cowplot 1.2.0 (89) packages, and repetitive elements were colored using the colorRampPalette feature on the “Paired” palette in RColorBrewer v.1.1-3. Treemap tile figures were visualized using the Treemapify v2.5.6 package (90), and the size of each treemap figure was scaled based on the total abundance of its TEs compared to the one with the highest TE abundance. For the analysis of the intra-variation of centromeric region sizes we used the newly generated IMI230595m assembly for *P. pseudocomata* and high-quality genome assemblies for 11 *P. anserina* strains (Supplementary Table 2). We located the centromere flanking orthologous genes from the PaWa63p (*P. anserina*) and CBS415.72m (*P. pseudocomata*) strains, respectively, and by using IGV v2.14.1 (85).

### Estimation of the divergence time of the *P. anserina* species complex

To estimate the divergence time of the *P. anserina* species complex we assumed a simple neutral model with mutation-drift equilibrium, infinite-sites, and free-recombination (following (91)). We calculated the synonymous nucleotide divergence (*d_S_*) between *P. anserina* and each of the other species, from previously-produced alignments of 100 genes randomly sampled from the genome (39). For each species pair, we estimated divergence in generations as *Tg* = *d_S_*/(2 × *k*), where *k* is the neutral substitution rate per site (92). As *P. anserina* can complete its life cycle in around 15 days in laboratory conditions, we assumed that there were around 365/11 = 24 generations per year. Hence, we estimated the divergence time in years as *Ty* = *Tg* / 24 (91). As the value of *k* is unknown for *Podospora,* we used the upper and lower bound estimates of Eurotiomycetes, corresponding to 0.9e-9 and 16.7e-9 substitution per site per year, respectively (40). We then took the median of the divergence time of all genes to get species-pairs divergence estimates for a given *k*. Finally, we averaged the median divergence times of the species-pairs comparison to get the age of the species complex. The sister species of *P. anserina*, *P. pauciseta*, was ignored for this calculation as their most recent common ancestor is necessarily younger than the species complex (30).

### Transposable element retrieval and subsequent analyses

To retrieve BLAST hit haplotypes of the TE families from the genomes of the seven species we used the representative sequences from the PodoTE-2.00.lib repeat library (33) and the script query2haplotype.py v1.92 (https://github.com/SLAment/Genomics) with the parameters –extrabp 500 –vicinity 1000 for retrotransposon families (*arthroleptis, rana, xenopus*). For the LTR-retrotransposons we only collected the LTRs instead of the full elements as this will capture both recent insertions and older ones that only contain solo-LTRs. For the DNA transposon *discoglosse* we instead used –extrabp 1000 –vicinity 5000, as this better captured the full copies of this element compared to the smaller LTRs. Collected copies were curated by first aligning the copies to the representative sequence to remove short fragmented sequences and split up haplotypes with multiple copies based on LAST dotplot alignments from the online MAFFT website (93). Next, the remaining copies were aligned again for a second round and oriented in the same orientation as the consensus. The alignment was then trimmed to remove the flanking regions outside the ends of the element and then finally poorly aligning copies still remaining were removed. For the *crapaud* LTR family phylogeny, we modified the LTR phylogeny from (33) to also include whether each copy was found in a centromeric region or not.

To compare copy sequence divergences from the representative sequences in *P. pseudocomata* we used the calculated Kimura2-parameter TE-family average values from the RepeatMasker script calcDivergenceFromAlign.pl. To identify RIPped copies in *P. pseudocomata* we retrieved a multi-fasta file of copies identified by RepeatMasker, which we then used as input in TheRIPper. To get a single RIP estimate for all copies, we set the window size to 10000 and proceeded with the default setting for all other parameters. We used the RIP composite index, where the threshold for a sequence being RIPped is 0, i.e., anything above zero is considered to be RIP affected.

### Permutation test and transposable element maximum-likelihood phylogenies

To test associations between transposable elements and centromeric regions in each of the species we divided the genomes into 20kb long non-overlapping windows and calculated the GC-content, proportion of genes, and proportion of masked TE bp in each window. Since the genomes of the species complex are gene dense we filtered the windows to only compare the centromeric regions with repeat rich non-centromeric windows, *i.e.,* windows within the top ten percentiles in regards to TE-content. Permutation tests were carried out using the permTest function in the R package RegioneR v1.34.0 (94) with 1000 permutations, the resampleRegions randomization function, and counting the number of overlaps between the centromeric regions and other repeat rich windows.

Maximum likelihood phylogenies of the *discoglosse* TE were generated using IQ-TREE v2.2.2.6 (95) using ultra fast bootstrap support using 1000 replicates (96) and single blanch support SH-aLRT using 1000 replicates (97). The GTR+F+R8 model was used as suggested by ModelFinder (98). For the *crapaud* TE we used the terminal repeat maximum-likelihood phylogeny from (33). For *arthroleptis*, *rana*, and *xenopus* we used MSA alignments of the terminal repeats to generate phylogenies using IQ-TREE v 1.6.12 (99) with ultra fast bootstrap (1000 replicates) and SH-aLRT (1000 replicates). The models used for the three TE families (in the same order as above) based on ModelFinder were: K2P+G4, TNe+G4, and K2P+G4. For visualization we used the interactive tree of life website (100) for *discoglosse* and *crapaud;* and FigTree v1.4.4 (101) for *arthroleptis*, *rana*, and *xenopus*.

## Data availability

Reads and genome assembly will be deposited in ENA. Assembly annotation and other data files can be found on Figshare (https://doi.org/10.6084/m9.figshare.30636149). Scripts and accompanying data can be found on GitHub (https://github.com/Ivwster/Podospora_Centromeres/).

## Contributions

IW and HJ initiated the study. ML, IW, EM, LS and PG performed the wet lab experiments. IW performed the bioinformatic analyses, with the exception of the *P. anserina* divergence time estimation, which was performed by SLAV. PS acquired the strain IMI230595 from CABI Genetic Resource Collection, reidentified it as *P. pseudocomata*, and provided F1 homokaryotic progeny for this strain. AV sequenced the genome of IMI230595 and SLAV assembled and annotated it. IW drafted the manuscript with contributions from all other authors. IW, AV, PG, FM, and HJ contributed with conceptualization of the study. All authors contributed with interpretation of results. HJ supervised the study. HJ, AV, and FM contributed with necessary resources for the completion of the study. All authors read and approved the final manuscript.

## Acknowledgements and Funding

We acknowledge the CABI Genetic Resource Collection for providing the culture for the strain IMI230595.We acknowledge the Swedish National Genomics Infrastructure (NGI) for support on Pacbio sequencing of IMI230595m and for ChIP-seq library preparation and sequencing of *P. pseudocomata* samples. We acknowledge Novogene for ChIP-seq library preparation and sequencing of *P. anserina* samples.

The computations/data handling was enabled by resources provided by the National Academic Infrastructure for Supercomputing in Sweden (NAISS), partially funded by the Swedish Research Council through grant agreement no. 2022-06725.

IW and HJ contributions were funded by the Bergianus Foundation at the Royal Swedish Academy of Sciences. PG, ML, EM and FM contribution was funded by the Diversity of Biological Mechanisms Programme of the Institut National des Sciences Biologiques (INSB) du Centre National de la Recherche Scientifique (CNRS). SLAV was supported by the Swedish Research Council (grant 2022-00341). LS was supported by the Carl Trygger Foundation (project CTS 20:477 to AV). AV was supported by the Swedish Research Council (grant 2021-04290) and by Lars Hiertas Minne and Nilsson-Ehle endowment for the sequencing.

## Notes

https://doi.org/10.6084/m9.figshare.30636149

https://github.com/Ivwster/Podospora_Centromeres/

### Competing Interest Statement

The authors have declared no competing interest.

## References

1. G. Bourque, et al., Ten things you should know about transposable elements. Genome Biol. 19, 199 (2018).

2. N. Buchon, C. Vaury, RNAi: a defensive RNA-silencing against viruses and transposable elements. Heredity 96, 195–202 (2006).

3. E. Gladyshev, Repeat-Induced Point Mutation (RIP) and Other Genome Defense Mechanisms in Fungi. Microbiol. Spectr. 5 (2017).

4. R. L. Cosby, N.-C. Chang, C. Feschotte, Host–transposon interactions: conflict, cooperation, and cooption. Genes Dev. 33, 1098–1116 (2019).

5. E. U. Selker, E. B. Cambareri, B. C. Jensen, K. R. Haack, Rearrangement of duplicated DNA in specialized cells of Neurospora. Cell 51, 741–752 (1987).

6. E. B. Cambareri, B. C. Jensen, E. Schabtach, E. U. Selker, Repeat-induced G-C to A-T mutations in Neurospora. Science 244, 1571–1575 (1989).

7. E. B. Cambareri, M. J. Singer, E. U. Selker, Recurrence of repeat-induced point mutation (RIP) in Neurospora crassa. Genetics 127, 699–710 (1991).

8. E. Gladyshev, N. Kleckner, Direct recognition of homology between double helices of DNA in Neurospora crassa. Nat. Commun. 5, 3509 (2014).

9. E. Gladyshev, N. Kleckner, Recombination-independent recognition of DNA homology for repeat-induced point mutation. Curr. Genet. 63, 389–400 (2017).

10. A. K. Mazur, E. Gladyshev, Partition of Repeat-Induced Point Mutations Reveals Structural Aspects of Homologous DNA-DNA Pairing. Biophys. J. 115, 605–615 (2018).

11. K. Guin, L. Sreekumar, K. Sanyal, Implications of the Evolutionary Trajectory of Centromeres in the Fungal Kingdom. Annu. Rev. Microbiol. 74, 835–853 (2020).

12. P. B. Talbert, S. Henikoff, What makes a centromere? Exp. Cell Res. 389, 111895 (2020).

13. S. Henikoff, K. Ahmad, H. S. Malik, The centromere paradox: stable inheritance with rapidly evolving DNA. Science 293, 1098–1102 (2001).

14. S. G. Nergadze, et al., Birth, evolution, and transmission of satellite-free mammalian centromeric domains. Genome Res. 28, 789–799 (2018).

15. F. M. Piras, et al., Molecular Dynamics and Evolution of Centromeres in the Genus Equus. Int. J. Mol. Sci. 23, 4183 (2022).

16. G. A. Logsdon, et al., The variation and evolution of complete human centromeres. Nature 1–10 (2024). 10.1038/s41586-024-07278-3.

17. L. C. Oliveira, G. A. Torres, Plant centromeres: genetics, epigenetics and evolution. Mol. Biol. Rep. 45, 1491–1497 (2018).

18. M. Naish, et al., The genetic and epigenetic landscape of the Arabidopsis centromeres. Science 374, eabi7489 (2021).

19. M. Naish, I. R. Henderson, The structure, function, and evolution of plant centromeres. Genome Res. 34, 161–178 (2024).

20. K. M. Smith, P. A. Phatale, C. M. Sullivan, K. R. Pomraning, M. Freitag, Heterochromatin Is Required for Normal Distribution of Neurospora crassa CenH3. Mol. Cell. Biol. 31, 2528–2542 (2011).

21. V. Yadav, et al., RNAi is a critical determinant of centromere evolution in closely related fungi. Proc. Natl. Acad. Sci. 115, 3108–3113 (2018).

22. P. Silar, Podospora anserina (2020). 978-2-9555841-2-5. ⟨hal-02475488⟩

23. P. Silar, et al., The Podospora anserina species complex in metropolitan and overseas France with description of a new species, Podospora reunionensis sp. nov. Cryptogam. Mycol. (in press).

24. P. Silar, “Podospora anserina: From Laboratory to Biotechnology” in Genomics of Soil-and Plant-Associated Fungi, Soil Biology., B. A. Horwitz, P. K. Mukherjee, M. Mukherjee, C. P. Kubicek, Eds. (Springer, 2013), pp. 283–309.

25. F. Graïa, et al., Genome quality control: RIP (repeat-induced point mutation) comes to Podospora. Mol. Microbiol. 40, 586–595 (2001).

26. K. Bouhouche, D. Zickler, R. Debuchy, S. Arnaise, Altering a gene involved in nuclear distribution increases the repeat-induced point mutation process in the fungus Podospora anserina. Genetics 167, 151–159 (2004).

27. P. Grognet, et al., A RID-like putative cytosine methyltransferase homologue controls sexual development in the fungus Podospora anserina. PLoS Genet. 15, e1008086 (2019).

28. A. A. Vogan, et al., Combinations of Spok genes create multiple meiotic drivers in Podospora. eLife 8, e46454 (2019).

29. A. A. Vogan, et al., The Enterprise, a massive transposon carrying Spok meiotic drive genes. Genome Res. 31, 789–798 (2021).

30. S. L. Ament-Velásquez, et al., High-Quality Genome Assemblies of 4 Members of the Podospora anserina Species Complex. Genome Biol. Evol. 16, evae034 (2024).

31. A. Hamann, F. Feller, H. D. Osiewacz, Yeti– a degenerate gypsy-like LTR retrotransposon in the filamentous ascomycete Podospora anserina. Curr. Genet. 38, 132–140 (2000).

32. E. Espagne, et al., The genome sequence of the model ascomycete fungus Podospora anserina. Genome Biol. 9, R77 (2008).

33. I. Westerberg, S. L. Ament-Velásquez, A. A. Vogan, H. Johannesson, Evolutionary dynamics of the LTR-retrotransposon crapaud in the Podospora anserina species complex and the interaction with repeat-induced point mutations. Mob. DNA 15, 1–16 (2024).

34. T. Meerupati, et al., Genomic Mechanisms Accounting for the Adaptation to Parasitism in Nematode-Trapping Fungi. PLOS Genet. 9, e1003909 (2013).

35. J. K. Hane, A. H. Williams, A. P. Taranto, P. S. Solomon, R. P. Oliver, “Repeat-Induced Point Mutation: A Fungal-Specific, Endogenous Mutagenesis Process” in Genetic Transformation Systems in Fungi, Volume 2, M. A. van den Berg, K. Maruthachalam, Eds. (Springer International Publishing, 2015), pp. 55–68.

36. M. Freitag, R. L. Williams, G. O. Kothe, E. U. Selker, A cytosine methyltransferase homologue is essential for repeat-induced point mutation in Neurospora crassa. Proc. Natl. Acad. Sci. U. S. A. 99, 8802–8807 (2002).

37. D. Marcou, A. Masson, J.-M. Simonet, G. Piquepaille, Evidence for Non-Random Spatial Distribution of Meiotic Exchanges in Podospora anserina: Comparison between Linkage Groups 1 and 6. Mol. Genet. Genomics MGG 176, 67–79 (1979).

38. C. Boucher, N. Tinh-Suong, P. Silar, Species Delimitation in the Podospora anserina/ P. pauciseta/P. comata Species Complex (Sordariales). Cryptogam. Mycol. 38, 485–506 (2017).

39. S. L. Ament-Velásquez, S. J. Saupe, NOD-like receptor genes evolve under diversity-enhancing mechanisms in a fungal species complex. [Preprint] (2025). Available at: https://www.biorxiv.org/content/10.1101/2025.09.29.679196v1 [Accessed 20 October 2025].

40. T. Kasuga, T. J. White, J. W. Taylor, Estimation of Nucleotide Substitution Rates in Eurotiomycete Fungi. Mol. Biol. Evol. 19, 2318–2324 (2002).

41. C. A. D’Souza, et al., Genome Variation in Cryptococcus gattii, an Emerging Pathogen of Immunocompetent Hosts. mBio 2, 10.1128/mbio.00342-10 (2011).

42. S. van Wyk, B. D. Wingfield, L. De Vos, N. A. van der Merwe, E. T. Steenkamp, Genome-Wide Analyses of Repeat-Induced Point Mutations in the Ascomycota. Front. Microbiol. 11, 622368 (2021).

43. T. Badet, D. Croll, Phylogenomic signatures of repeat-induced point mutations across the fungal kingdom. PLOS Biol. 23, e3003433 (2025).

44. L. Le Chevanton, G. Leblon, S. Lebilcot, Duplications created by transformation in Sordaria macrospora are not inactivated during meiosis. Mol. Gen. Genet. MGG 218, 390–396 (1989).

45. M. Walz, U. Kück, Transformation of Sordaria macrospora to hygromycin B resistance: characterization of transformants by electrophoretic karyotyping and tetrad analysis. Curr. Genet. 29, 88–95 (1995).

46. M. Nowrousian, et al., De novo Assembly of a 40 Mb Eukaryotic Genome from Short Sequence Reads: Sordaria macrospora, a Model Organism for Fungal Morphogenesis. PLoS Genet. 6, e1000891 (2010).

47. H. P. Cam, K. Noma, H. Ebina, H. L. Levin, S. I. S. Grewal, Host genome surveillance for retrotransposons by transposon-derived proteins. Nature 451, 431–436 (2008).

48. J. C. Matasyoh, B. Dittrich, A. Schueffler, H. Laatsch, Larvicidal activity of metabolites from the endophytic Podospora sp. against the malaria vector Anopheles gambiae. Parasitol. Res. 108, 561–566 (2011).

49. B. J. Chabot, et al., Transcription of a centromere-enriched retroelement and local retention of its RNA are significant features of the CENP-A chromatin landscape. Genome Biol. 25, 295 (2024).

50. A. Shimada, et al., Retrotransposon addiction promotes centromere function via epigenetically activated small RNAs. Nat. Plants 10, 1304–1316 (2024).

51. S. Dutta, K. Bhat, R. Aggarwal, K. Sanyal, Fungi as models of centromere innovation: from DNA sequence to 3-dimensional arrangement. Chromosome Res. 33, 18 (2025).

52. G. C. Ferreri, et al., Recent Amplification of the Kangaroo Endogenous Retrovirus, KERV, Limited to the Centromere. J. Virol. 85, 4761–4771 (2011).

53. D. Gao, N. Jiang, R. A. Wing, J. Jiang, S. A. Jackson, Transposons play an important role in the evolution and diversification of centromeres among closely related species. Front. Plant Sci. 6, 216 (2015).

54. C.-H. Chang, et al., Islands of retroelements are major components of Drosophila centromeres. PLOS Biol. 17, e3000241 (2019).

55. M. F. Seidl, et al., Repetitive Elements Contribute to the Diversity and Evolution of Centromeres in the Fungal Genus Verticillium. mBio 11, e01714–20 (2020).

56. P. Wlodzimierz, et al., Cycles of satellite and transposon evolution in Arabidopsis centromeres. Nature 618, 557–565 (2023).

57. T. Sultana, A. Zamborlini, G. Cristofari, P. Lesage, Integration site selection by retroviruses and transposable elements in eukaryotes. Nat. Rev. Genet. 18, 292–308 (2017).

58. H. S. Malik, T. H. Eickbush, Modular Evolution of the Integrase Domain in the Ty3/Gypsy Class of LTR Retrotransposons. J. Virol. 73, 5186–5190 (1999).

59. B. Gorinšek, F. Gubenšek, D. Kordiš, Evolutionary Genomics of Chromoviruses in Eukaryotes. Mol. Biol. Evol. 21, 781–798 (2004).

60. X. Gao, Y. Hou, H. Ebina, H. L. Levin, D. F. Voytas, Chromodomains direct integration of retrotransposons to heterochromatin. Genome Res. 18, 359–369 (2008).

61. A. C. Spradling, H. J. Bellen, R. A. Hoskins, Drosophila P elements preferentially transpose to replication origins. Proc. Natl. Acad. Sci. 108, 15948–15953 (2011).

62. M. J. Curcio, S. Lutz, P. Lesage, The Ty1 LTR-Retrotransposon of Budding Yeast, Saccharomyces cerevisiae. Microbiol. Spectr. 3, 10.1128/microbiolspec.mdna3-0053-2014 (2015).

63. G. Rizet, C. Engelman, Contribution à l’étude génétique d’un ascomycete tétrasporé: Podospora anserina. Rev Cytol Biol Végétale (11), 201–304. (1949).

64. M. van der Gaag, et al., Spore-Killing Meiotic Drive Factors in a Natural Population of the Fungus Podospora anserina. Genetics 156, 593–605 (2000).

65. Y. Sun, J. Svedberg, M. Hiltunen, P. Corcoran, H. Johannesson, Large-scale suppression of recombination predates genomic rearrangements in Neurospora tetrasperma. Nat. Commun. 8, 1140 (2017).

66. H. Cheng, G. T. Concepcion, X. Feng, H. Zhang, H. Li, Haplotype-resolved de novo assembly using phased assembly graphs with hifiasm. Nat. Methods 18, 170–175 (2021).

67. S. Kurtz, et al., Versatile and open software for comparing large genomes. Genome Biol. 5, R12 (2004).

68. C. Holt, M. Yandell, MAKER2: an annotation pipeline and genome-database management tool for second-generation genome projects. BMC Bioinformatics 12, 491 (2011).

69. M. S. Campbell, C. Holt, B. Moore, M. Yandell, Genome Annotation and Curation Using MAKER and MAKER-P. Curr. Protoc. Bioinforma. 48, 4.11.1–4.11.39 (2014).

70. G. Lelandais, D. Remy, F. Malagnac, P. Grognet, New insights into genome annotation in Podospora anserina through re-exploiting multiple RNA-seq data. BMC Genomics 23, 859 (2022).

71. P. Silar, et al., A gene graveyard in the genome of the fungus Podospora comata. Mol. Genet. Genomics MGG 294, 177–190 (2019).

72. J. M. Palmer, J. Stajich, Funannotate v1.8.1: Eukaryotic genome annotation. (2020). 10.5281/zenodo.4054262. Deposited 28 September 2020.

73. P. Silar, Two new easy to use vectors for transformations. Fungal Genet. Rep. 42, 73 (1995).

74. Y. Brygoo, R. Debuchy, Transformation by integration in Podospora anserina. Mol. Gen. Genet. MGG 200, 128–131 (1985).

75. F. Carlier, et al., Loss of EZH2-like or SU(VAR)3–9-like proteins causes simultaneous perturbations in H3K27 and H3K9 tri-methylation and associated developmental defects in the fungus Podospora anserina. Epigenetics Chromatin 14, 22 (2021).

76. S. Chen, Ultrafast one-pass FASTQ data preprocessing, quality control, and deduplication using fastp. iMeta 2, e107 (2023).

77. A. M. Bolger, M. Lohse, B. Usadel, Trimmomatic: a flexible trimmer for Illumina sequence data. Bioinforma. Oxf. Engl. 30, 2114–2120 (2014).

78. H. Li, R. Durbin, Fast and accurate long-read alignment with Burrows-Wheeler transform. Bioinforma. Oxf. Engl. 26, 589–595 (2010).

79. H. Li, et al., The Sequence Alignment/Map format and SAMtools. Bioinforma. Oxf. Engl. 25, 2078–2079 (2009).

80. F. Ramírez, et al., deepTools2: a next generation web server for deep-sequencing data analysis. Nucleic Acids Res. 44, W160–W165 (2016).

81. Y. Zhang, et al., Model-based Analysis of ChIP-Seq (MACS). Genome Biol. 9, R137 (2008).

82. A. Smit, R. Hubley, P. Green, RepeatMasker Open-4.0 (1996–2015). Available at: https://www.repeatmasker.org

83. L. Lopez-Delisle, et al., pyGenomeTracks: reproducible plots for multivariate genomic datasets. Bioinformatics 37, 422–423 (2021).

84. S. van Wyk, et al., The RIPper, a web-based tool for genome-wide quantification of Repeat-Induced Point (RIP) mutations. PeerJ 7, e7447 (2019).

85. J. T. Robinson, et al., Integrative Genomics Viewer. Nat. Biotechnol. 29, 24–26 (2011).

86. A. L. Delcher, Fast algorithms for large-scale genome alignment and comparison. Nucleic Acids Res. 30, 2478–2483 (2002).

87. R Core Team. R: A Language and Environment for Statistical Computing. R Foundation for Statistical Computing, Vienna, Austria (2021).

88. H. Wickham, ggplot2: Elegant Graphics for Data Analysis (Springer-Verlag New York, 2016).

89. C. O. Wilke, cowplot: Streamlined plot theme and plot annotations for ggplot2. Comprehensive R Archive Network (2025). 10.32614/CRAN.package.cowplot

90. D. Wilkins, treemapify: Draw treemaps in ggplot2. Comprehensive R Archive Network (2023). 10.32614/CRAN.package.treemapify

91. L. Guyot, et al., Sheltered load in fungal mating-type chromosomes revealed by fitness experiments. J. Evol. Biol. 38, 1256–1271 (2025).

92. John H. Gillespie, Population genetics: A concise guide, Second ed. (The Johns Hopkins University Press, 2004).

93. K. Katoh, J. Rozewicki, K. D. Yamada, MAFFT online service: multiple sequence alignment, interactive sequence choice and visualization. Brief. Bioinform. 20, 1160–1166 (2019).

94. B. Gel, et al., regioneR: an R/Bioconductor package for the association analysis of genomic regions based on permutation tests. Bioinforma. Oxf. Engl. 32, 289–291 (2016).

95. B. Q. Minh, et al., IQ-TREE 2: New Models and Efficient Methods for Phylogenetic Inference in the Genomic Era. Mol. Biol. Evol. 37, 1530–1534 (2020).

96. D. T. Hoang, O. Chernomor, A. von Haeseler, B. Q. Minh, L. S. Vinh, UFBoot2: Improving the Ultrafast Bootstrap Approximation. Mol. Biol. Evol. 35, 518–522 (2018).

97. S. Guindon, et al., New Algorithms and Methods to Estimate Maximum-Likelihood Phylogenies: Assessing the Performance of PhyML 3.0. Syst. Biol. 59, 307–321 (2010).

98. S. Kalyaanamoorthy, B. Q. Minh, T. K. F. Wong, A. von Haeseler, L. S. Jermiin, ModelFinder: fast model selection for accurate phylogenetic estimates. Nat. Methods 14, 587–589 (2017).

99. L.-T. Nguyen, H. A. Schmidt, A. von Haeseler, B. Q. Minh, IQ-TREE: A Fast and Effective Stochastic Algorithm for Estimating Maximum-Likelihood Phylogenies. Mol. Biol. Evol. 32, 268–274 (2015).

100. I. Letunic, P. Bork, Interactive Tree Of Life (iTOL): an online tool for phylogenetic tree display and annotation. Bioinformatics 23, 127–128 (2007).

101. A. Rambaut, FigTree, version 1.4.4 (2018). Available at: https://github.com/rambaut/figtree

