## Supplementary Figures for "A recent shift in centromere size and DNA content in *Podospora pseudocomata* co-occurs with the loss of a fungal genome defense system"

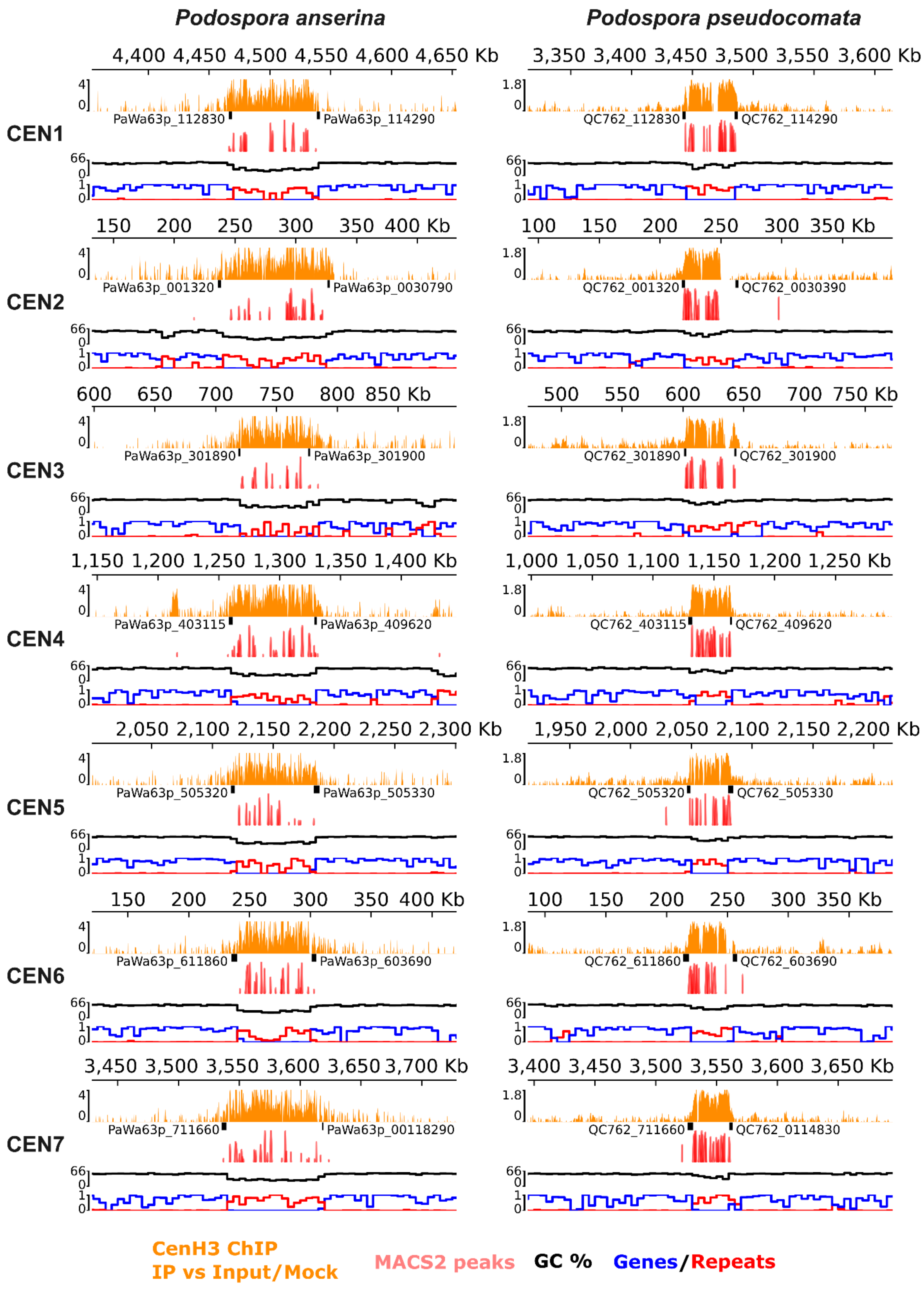

**Supplementary figure 1**: 300kb tracks centered at the mid-point of the cenH3 enrichment of the centromere regions of all seven chromosomes of both *P. anserina* and *P. pseudocomata*. Tracks from top to bottom: 1) Enrichment of immunoprecipitated cenH3 compared to input (*P. anserina*) or mock (*P. pseudocomata*). 2) The genes flanking the centromeric regions in black and their IDs in the respective assembly annotation. 3) Peaks called by MACS2 (q < 0.001). 4) GC % calculated in 5kb windows. 5) Gene (blue) and repeat (red) proportions calculated in 5kb non-overlapping windows.

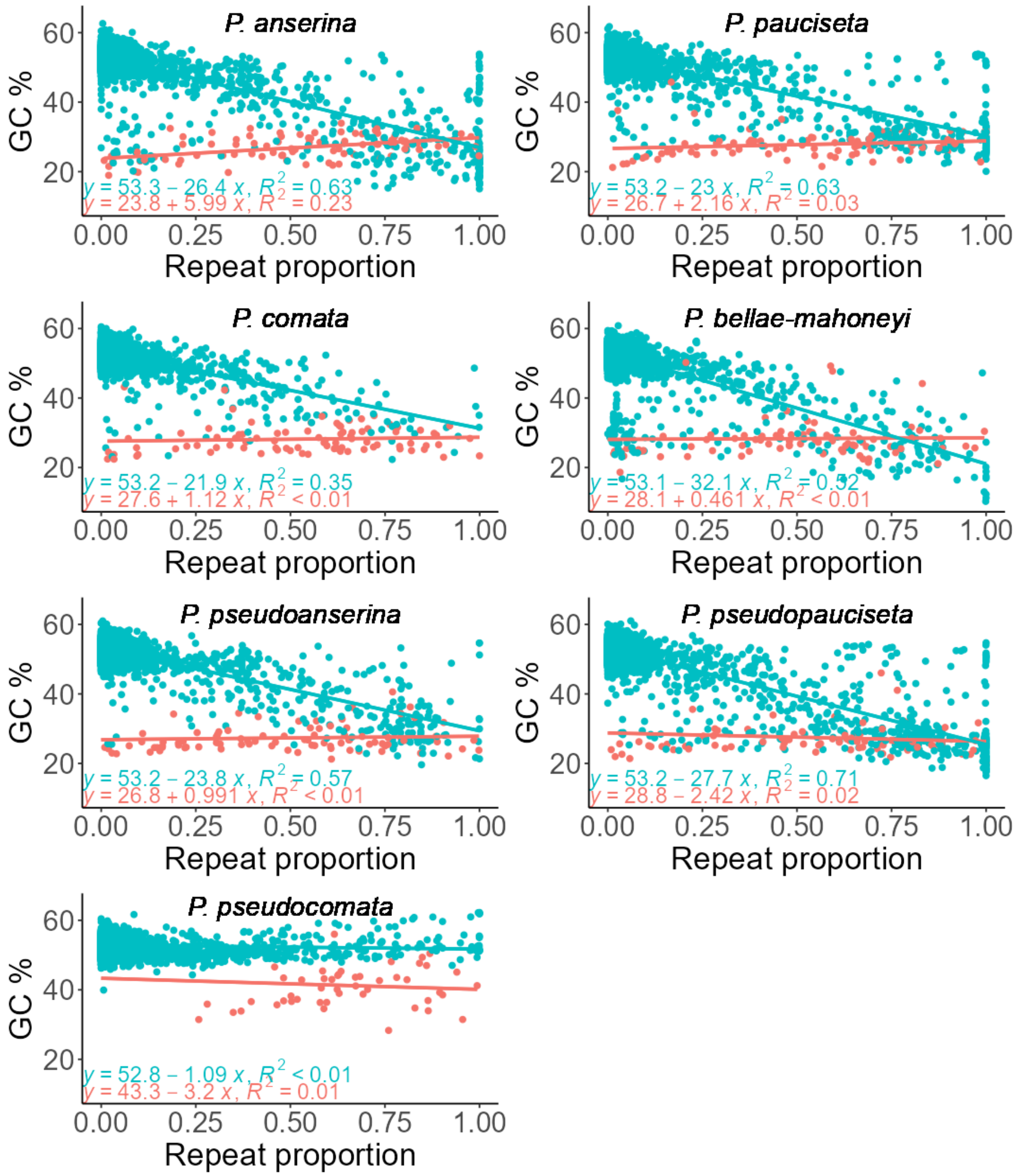

**Supplementary figure 2**: GC-content of centromeric 5kb non-overlapping sliding genomic windows of centromeric (red) and non-centromeric (blue) windows of the seven species in the *P. anserina* species complex

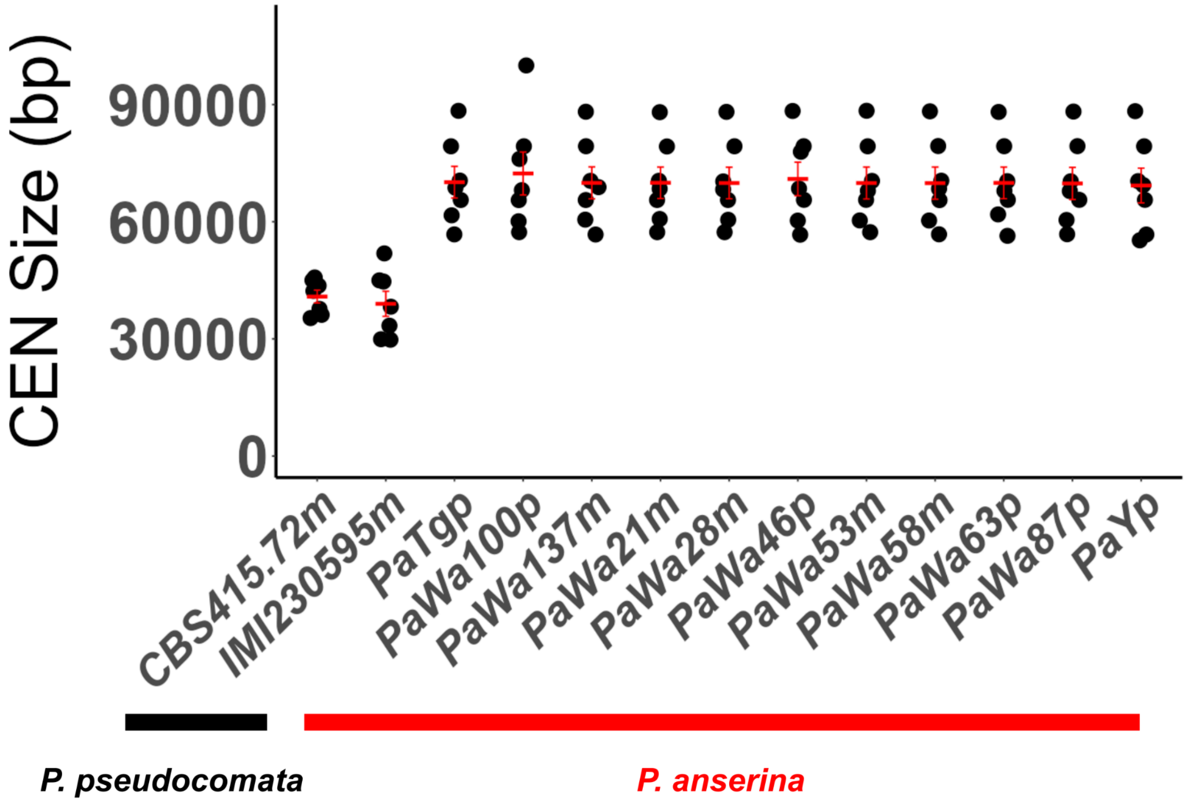


**Supplementary figure 3**: Intraspecies variation in centromeric region sizes in *P. pseudocomata* and *P. anserina* strains.

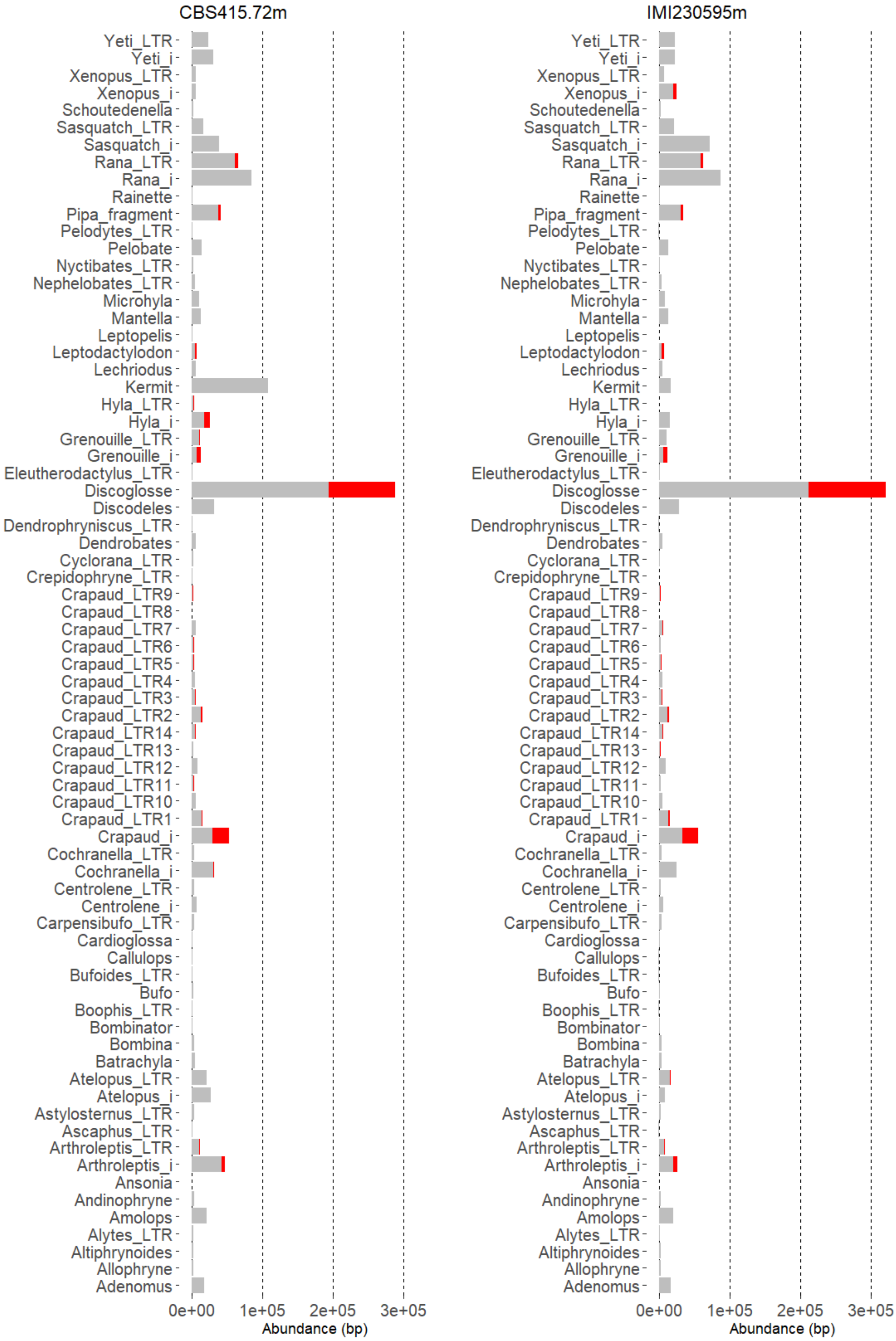


**Supplementary figure 4**: Transposable element abundance in base pairs in the two *Podospora pseudocomata* genomes CBS415.72m and newly generated IMI230595m in non-centromeric (grey) and centromeric (red) regions.

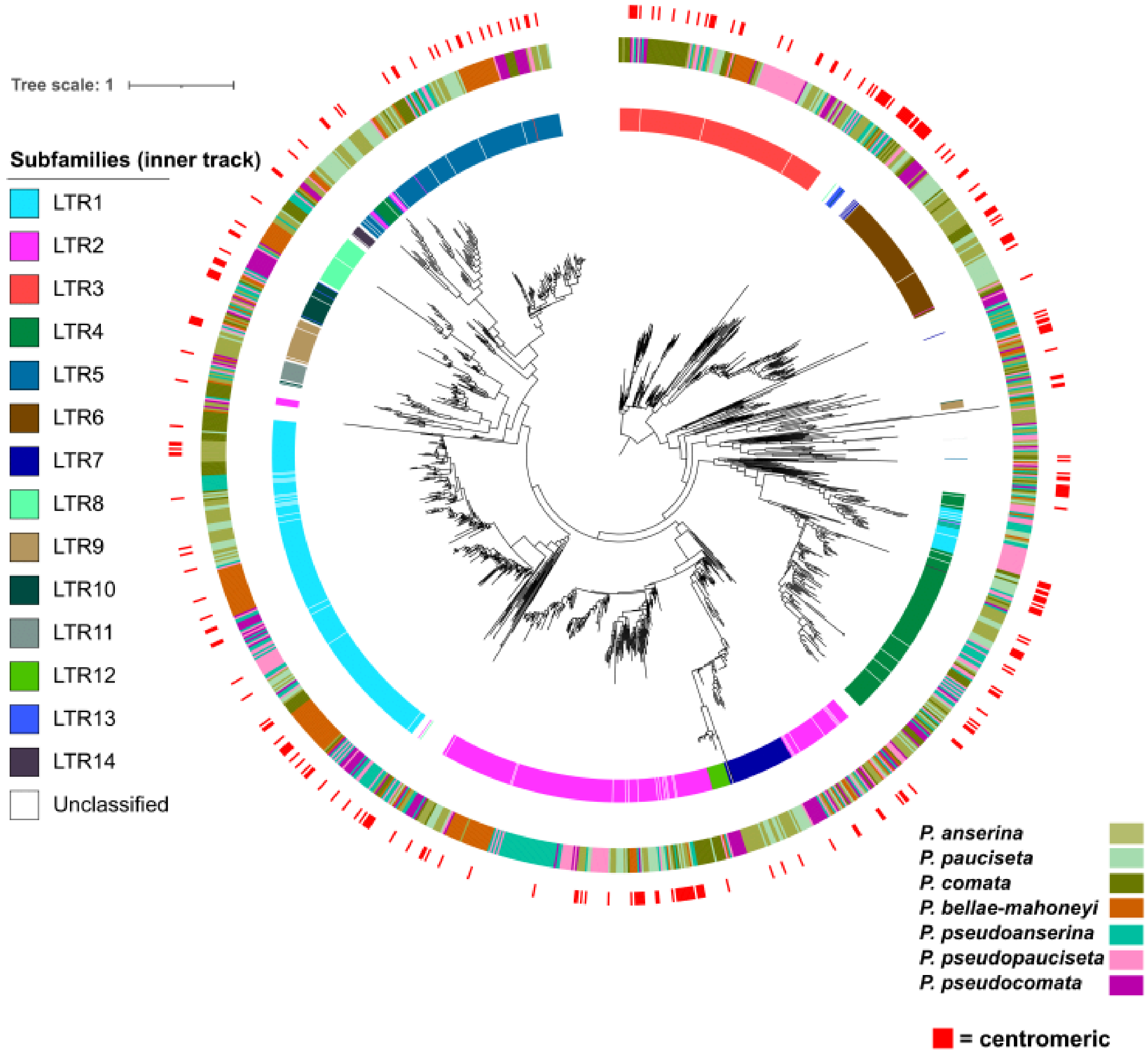


**Supplementary figure 5**: Phylogenetic tree adapted from Westerberg *et al.* 2024 showing copies of the different *crapaud* LTR subfamilies (inner ring), species (middle ring), and centromeric copies (outer ring).

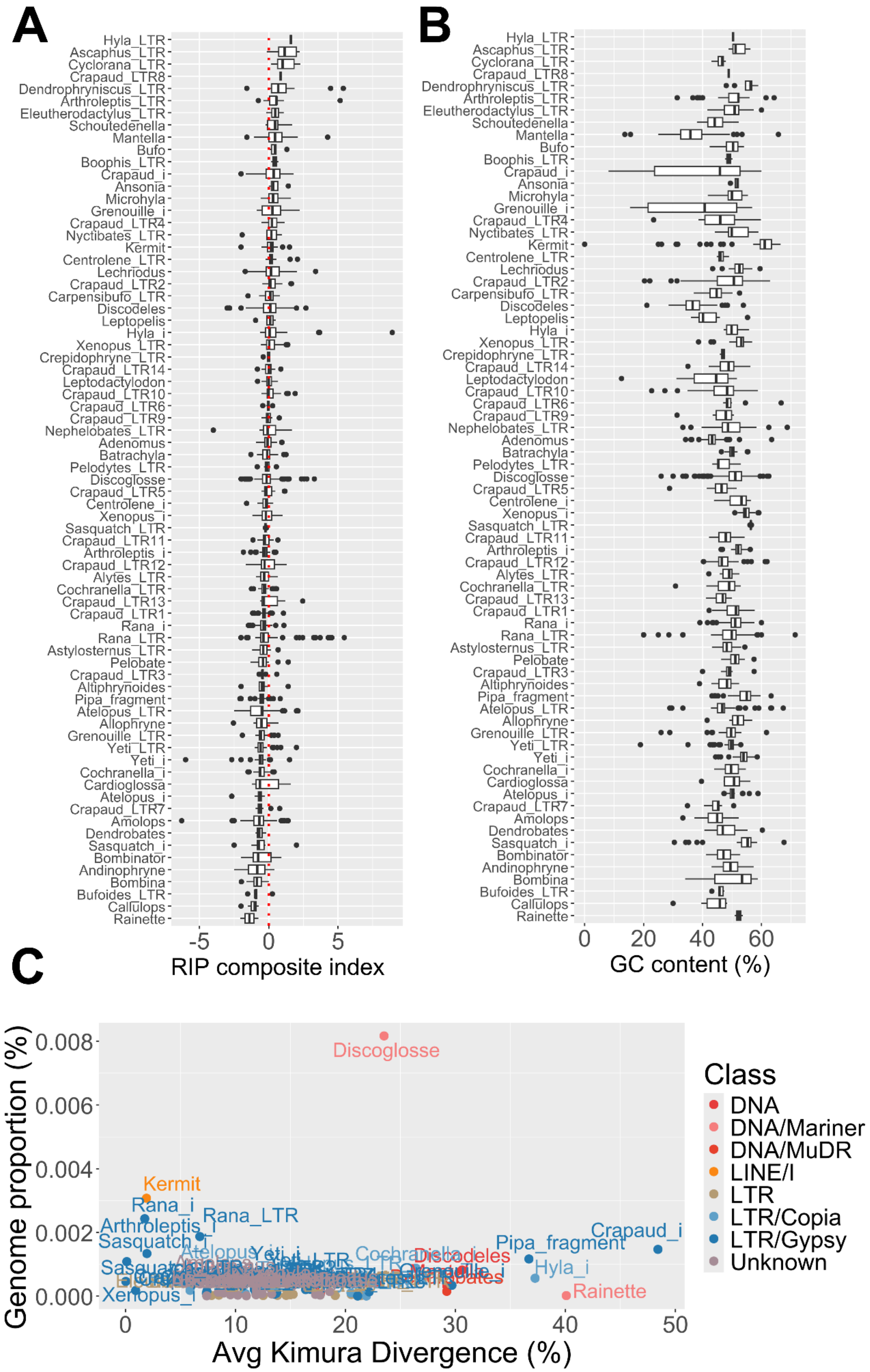
A
Hyla_LTR
Ascaphus_LTR
Cyclorana_LTR
Crapaud_LTR8
Dendrophryniscus_LTR
Arthroleptis_LTR
Eleutherodactylus_LTR
Schoutedenella
Mantella
Bufo
Boophis_LTR
Crapaud_i
Ansonia
Microhyla
Grenouille_i
Crapaud_LTR4
Nyctibates_LTR
Kermit
Centrolene_LTR
Lechriodus
Crapaud_LTR2
Carpensibufo_LTR
Discodeles
Leptopelis
Hyla_i
Xenopus_LTR
Crepidophryne_LTR
Crapaud_LTR14
Leptodactylodon
Crapaud_LTR10
Crapaud_LTR6
Crapaud_LTR9
Nephelobates_LTR
Adenomus
Batrachyla
Pelodytes_LTR
Discoglosse
Crapaud_LTR5
Centrolene_i
Xenopus_i
Sasquatch_LTR
Crapaud_LTR11
Arthroleptis_i
Crapaud_LTR12
Alytes_LTR
Cochranella_LTR
Crapaud_LTR13
Crapaud_LTR1
Rana_i
Rana_LTR
Astylosternus_LTR
Pelobate
Crapaud_LTR3
Altiphrynoides
Pipa_fragment
Atelopus_LTR
Allophryne
Grenouille_LTR
Yeti_LTR
Yeti_i
Cochranella_i
Cardioglossa
Atelopus_i
Crapaud_LTR7
Amolops
Dendrobates
Sasquatch_i
Bombinator
Andinophryne
Bombina
Bufoides_LTR
Callulops
Rainette
-5
0
5
RIP composite index
B
Hyla_LTR
Ascaphus_LTR
Cyclorana_LTR
Crapaud_LTR8
Dendrophryniscus_LTR
Arthroleptis_LTR
Eleutherodactylus_LTR
Schoutedenella
Mantella
Bufo
Boophis_LTR
Crapaud_i
Ansonia
Microhyla
Grenouille_i
Crapaud_LTR4
Nyctibates_LTR
Kermit
Centrolene_LTR
Lechriodus
Crapaud_LTR2
Carpensibufo_LTR
Discodeles
Leptopelis
Hyla_i
Xenopus_LTR
Crepidophryne_LTR
Crapaud_LTR14
Leptodactylodon
Crapaud_LTR10
Crapaud_LTR6
Crapaud_LTR9
Nephelobates_LTR
Adenomus
Batrachyla
Pelodytes_LTR
Discoglosse
Crapaud_LTR5
Centrolene_i
Xenopus_i
Sasquatch_LTR
Crapaud_LTR11
Arthroleptis_i
Crapaud_LTR12
Alytes_LTR
Cochranella_LTR
Crapaud_LTR13
Crapaud_LTR1
Rana_i
Rana_LTR
Astylosternus_LTR
Pelobate
Crapaud_LTR3
Altiphrynoides
Pipa_fragment
Atelopus_LTR
Allophryne
Grenouille_LTR
Yeti_LTR
Yeti_i
Cochranella_i
Cardioglossa
Atelopus_i
Crapaud_LTR7
Amolops
Dendrobates
Sasquatch_i
Bombinator
Andinophryne
Bombina
Bufoides_LTR
Callulops
Rainette
0
20
40
60
GC content (%)
C
Genome proportion (%)
0.008
0.006
0.004
0.002
0.000
Discoglosse
Kermit
Rana_i
Rana_LTR
Arthroleptis_i
Sasquatch
Crapaud_i
Pipa_fragment
Discodeles
Hyla_i
Rainette
Xenopus_i
0
10
20
30
40
50
Avg Kimura Divergence (%)
Class
DNA
DNA/Mariner
DNA/MuDR
LINE/I
LTR
LTR/Copia
LTR/Gypsy
Unknown

**Supplementary figure 6**: Transposable element family statistics in *P. pseudocomata*. A) Boxplots of RIP composite index values calculated by TheRIPper for copies identified by RepeatMasker masking. Copies above zero (dotted red line) are considered to be affected by RIP mutations (76). Note that LTR retrotransposons have been split into terminal repeats (LTR) and inner region (i) of the element, hence why *crapaud* is present multiple times due to it having subfamilies based on differences in the LTRs. B) Boxplots of GC content percentages of copies in each TE family. C) Average Kimura2 substitution divergence of copies to the representative sequence of each TE family and their overall genomic proportion.

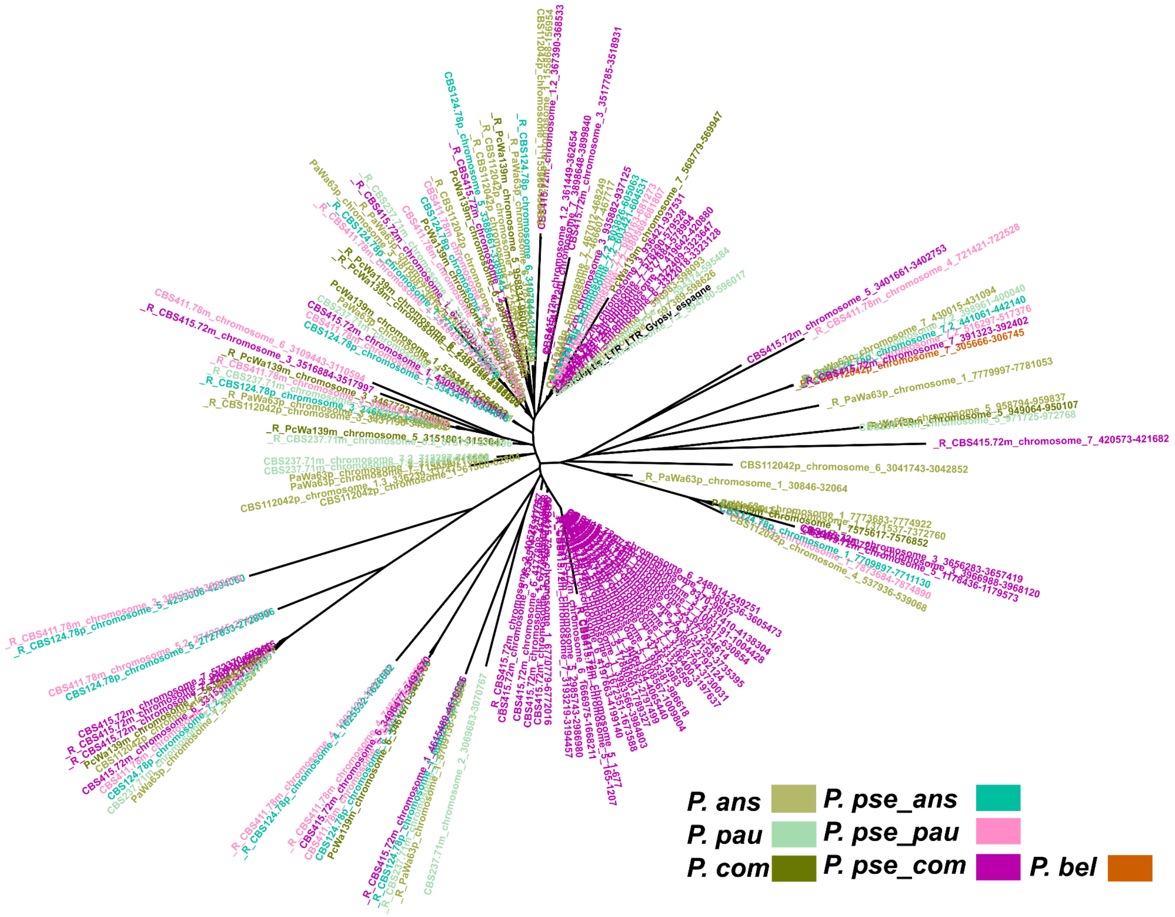


**Supplementary Figure 7**: Unrooted maximum likelihood phylogeny of terminal repeats from the *arthroleptis* LTR family. Tip names have been colored by species and the representative sequence is colored in black. *P. pse_ans* = *P. pseudoanserina*, *P. pse_pau* = *P. pseudopauciseta*, *P. pse_com* = *P. pseudocomata*.

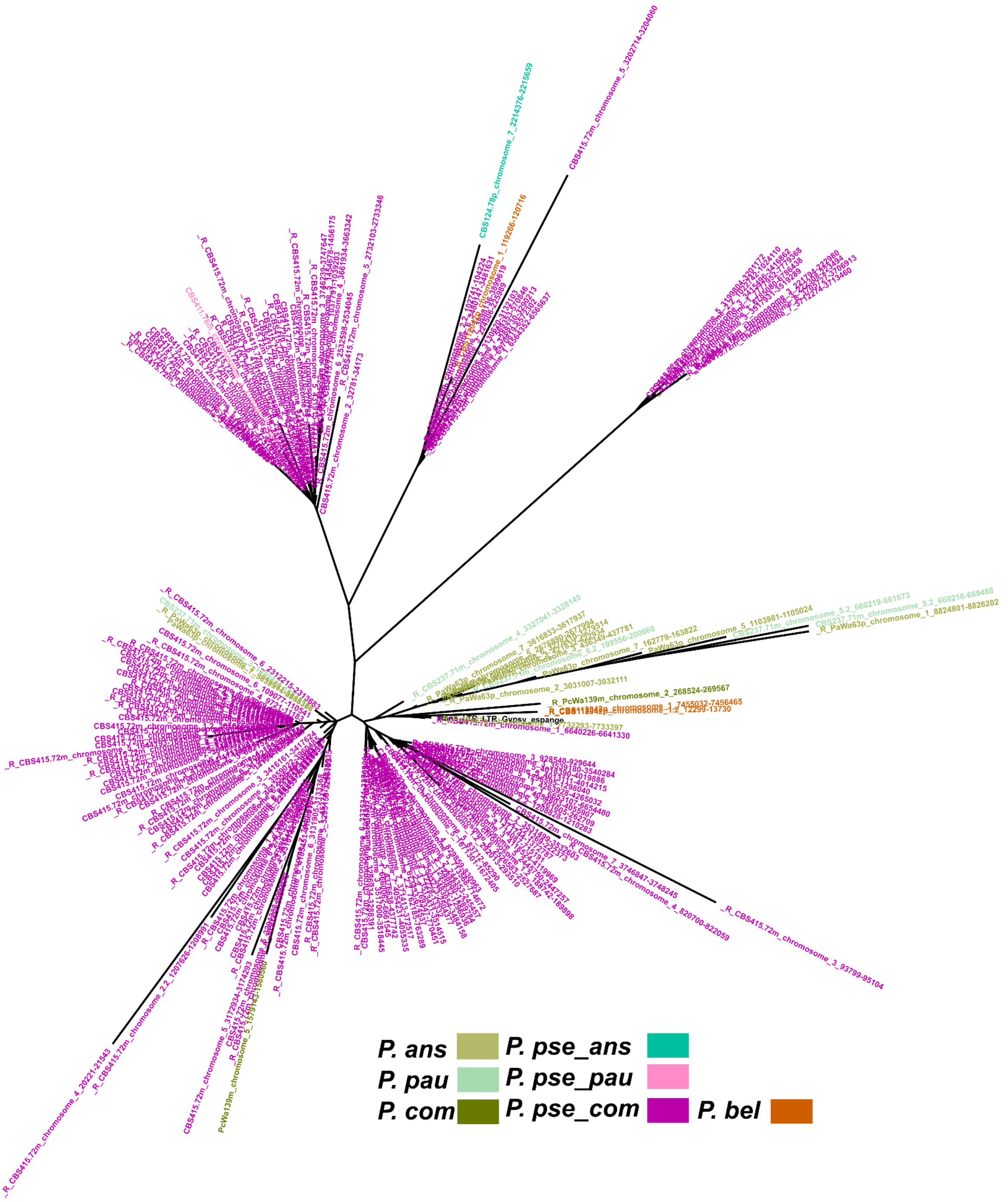


**Supplementary Figure 8**: Unrooted maximum likelihood phylogeny of terminal repeats from the *rana* LTR family. Tip names have been colored by species and the representative sequence is colored in black. *P. pse_ans* = *P. pseudoanserina*, *P. pse_pau* = *P. pseudopauciseta*, *P. pse_com* = *P. pseudocomata*.

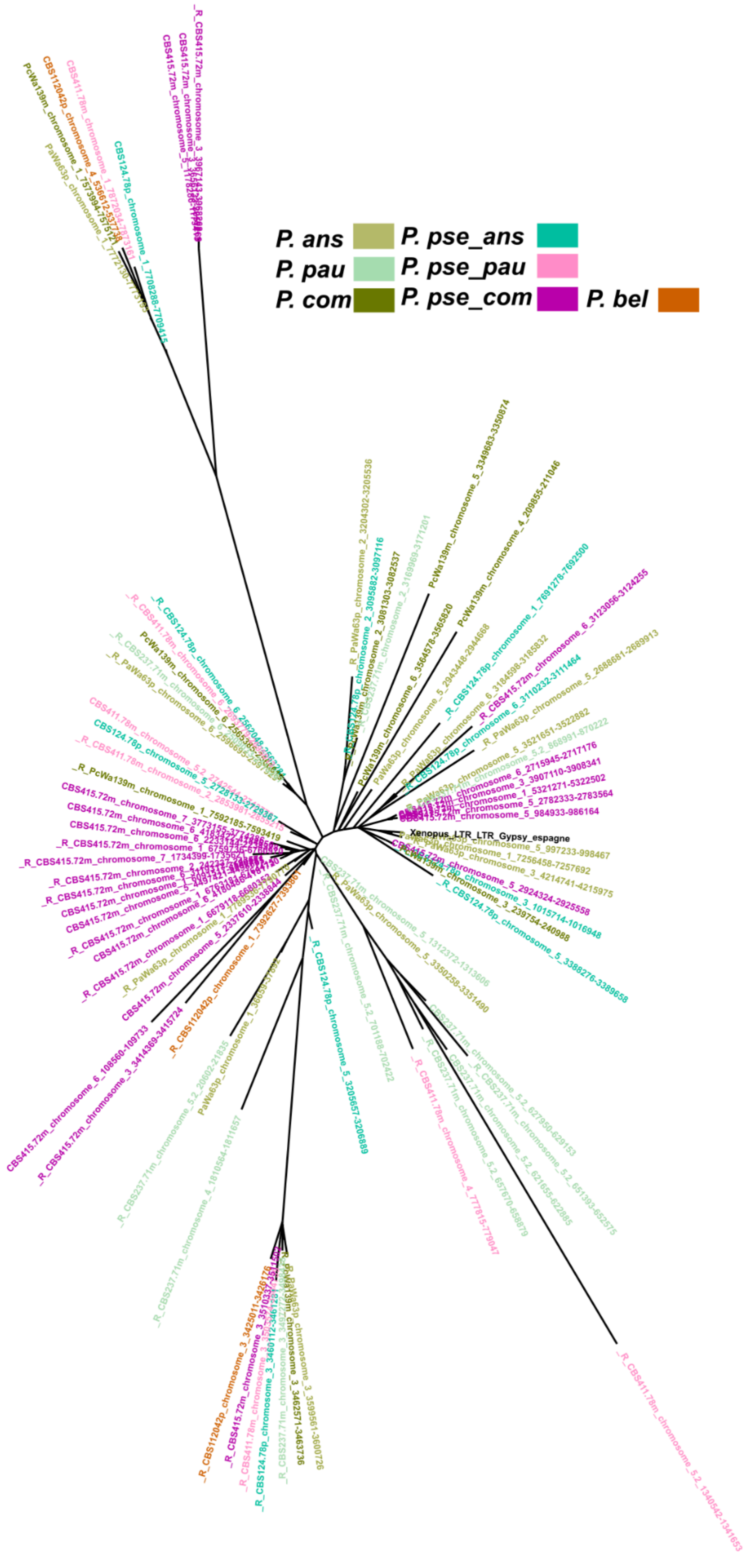


**Supplementary Figure 9**: Unrooted maximum likelihood phylogeny of terminal repeats from the *xenopus* LTR family. Tip names have been colored by species and the representative sequence is colored in black. *P. pse_ans* = *P. pseudoanserina*, *P. pse_pau* = *P. pseudopauciseta*, *P. pse_com* = *P. pseudocomata*.
